# A Predictive Coding Model of the N400

**DOI:** 10.1101/2023.04.10.536279

**Authors:** Samer Nour Eddine, Trevor Brothers, Lin Wang, Michael Spratling, Gina R. Kuperberg

## Abstract

The N400 event-related component has been widely used to investigate the neural mechanisms underlying real-time language comprehension. However, despite decades of research, there is still no unifying theory that can explain both its temporal dynamics and functional properties. In this work, we show that predictive coding – a biologically plausible algorithm for approximating Bayesian inference – offers a promising framework for characterizing the N400. Using an implemented predictive coding computational model, we demonstrate how the N400 can be formalized as the lexico-semantic prediction error produced as the brain infers meaning from linguistic form of incoming words. We show that the magnitude of lexico-semantic prediction error mirrors the functional sensitivity of the N400 to various lexical variables, priming, contextual effects, as well as their higher-order interactions. We further show that the dynamics of the predictive coding algorithm provide a natural explanation for the temporal dynamics of the N400, and a biologically plausible link to neural activity. Together, these findings directly situate the N400 within the broader context of predictive coding research, and suggest that the brain may use the same computational mechanism for inference across linguistic and non-linguistic domains.

## Introduction

A key discovery in the history of psycholinguistics was the presence of a neural signature of online language processing — the N400 event-related potential (ERP; Kutas & Hillyard, 1980, 1984). While there has been considerable interest in developing a theoretical framework for understanding the N400, this has proved a formidable challenge. Many theories and computational models have provided compelling explanations for its functional properties.

However, a unifying, biologically plausible account remains elusive. In this study, we show that the N400 can be understood as lexico-semantic prediction error, which is computed as a key step of *predictive coding* — a computational algorithm that has been proposed to carry out perceptual inference in the brain (Rao & Ballard, 1999; Mumford, 1992; Friston, 2005; Spratling, 2016b). Using an implemented predictive coding model of lexico-semantic processing, we show that the magnitude of lexico-semantic prediction error tracks the temporal dynamics of the N400 as well as its functional sensitivity to both bottom-up lexical and top-down contextual information.

The N400 ERP is a negative-going waveform that is detected at the scalp surface using both electroencephalography (EEG) and magnetoencephalography (MEG) between 300-500ms following the onset of any meaningful stimulus, such as a word or a picture (see Kutas & Federmeier, 2011, for a review). During language processing, the N400 is highly sensitive to the relationship between a word and its prior context, regardless of whether this context is a single word (in semantic and repetition priming paradigms, e.g. Bentin, McCarthy & Wood, 1985; Rugg, 1985), or a more extended sentence or discourse context (e.g., Kutas & Hillyard, 1984; DeLong, Urbach & Kutas, 2005; Van Berkum, Hagoort & Brown, 1999). The N400 is also elicited by words presented out of context where its amplitude is sensitive to several lexical variables, including orthographic neighborhood size (e.g. *core* > *kiwi*; Holcomb, Grainger & O’Rourke, 2002; Laszlo & Federmeier, 2011), lexical frequency (e.g. *wart* < *cold*; Rugg, 1990; Van Petten & Kutas, 1990), and concreteness/semantic richness (e.g. *lime* > *know*; Kounios & Holcomb, 1994; Holcomb, Kounios, Anderson & West, 1999; Rabovsky, Sommer & Abdel Rahman, 2012b).

Despite extensive work on the N400, there is still no general consensus on its functional significance. In the early 2000s, two competing theories dominated the debate: a lexical access and an integration account. Briefly, the lexical access account interpreted the N400 as reflecting the difficulty of accessing or “recognizing” a unique lexical item (e.g. Lau, Phillips, & Poeppel, 2008), while the integration account interpreted it as a “post-lexical” process that links the fully accessed item with its prior context (Brown & Hagoort, 1993; Hagoort, 2009). However, as several researchers pointed out, this type of dichotomy between “access” and “integration” has difficulty in explaining the sensitivity of the N400 to *both* lexical and contextual factors (Kutas & Federmeier, 2011; Baggio & Hagoort, 2011; Kuperberg 2016). More generally, this dichotomy rests on the somewhat questionable assumption that lexical access and semantic integration are distinct, separable cognitive processes that occur in a fixed sequence (see Laszlo & Federmeier, 2011; Kuperberg, Brothers & Wlotko, 2020 for discussion).

These shortcomings led to the more general proposal that the N400 reflects the impact of stimulus-driven activation on the current state of semantic memory (Kutas & Federmeier, 2011). In this framework, semantic memory is conceptualized as a dynamic multimodal system that is interactively influenced by both the high-level incremental interpretation of the prior context, as well as the linguistic form of each incoming word. This theory therefore intuitively explains a number of top-down and bottom-up influences on the amplitude of the N400. For example, as new bottom-up input activates overlapping orthographic neighbors, the co-activation of their semantic features would result in an enhanced N400 response to both words and non-words (see Laszlo & Federmeier, 2011), and the amplitude of the N400 would be reduced to the degree that the input matches semantic features had been pre-activated by the prior context, even if these inputs were lexically unexpected (e.g. Federmeier & Kutas, 1999). On the other hand, the theory’s flexibility leaves a number of cognitive mechanisms unspecified. How do particular stimuli activate the correct set of semantic features in long-term memory? Why does lexical processing result in the partial activation of orthographic and semantic neighbors, and how does the brain ultimately suppress these to settle on a “correct” interpretation of the bottom-up input? What determines the characteristic rise and fall of the N400 response? Most importantly, how are these processes implemented in a biologically plausible fashion in the brain?

One way of addressing these questions is through the development of explicit computational models. Several researchers have risen to this challenge, and a number of connectionist models of the N400 have been described (Laszlo & Plaut, 2012; Laszlo & Armstrong, 2014; Cheyette & Plaut, 2016; Rabovsky & McRae, 2014; Brouwer, Crocker, Venhuizen & Hoeks, 2017; Rabovsky, Hansen & McClelland, 2018; Rabovsky, 2020; Fitz & Chang, 2019), which we comprehensively review elsewhere (Nour Eddine, Brothers & Kuperberg, 2022). Broadly, these models of the N400 fall into two classes: word-level and sentence-level.

The word-level models were trained to map a single word-form input (e.g., a letter-string), clamped at the input layer, on to a pattern of activation that represented the word’s meaning at the top (output) layer (Laszlo & Plaut, 2012; Laszlo & Armstrong, 2014; Cheyette & Plaut, 2017; Rabovsky & McRae, 2014). In one set of studies, Laszlo, Plaut, Armstrong and Cheyette used a biologically motivated Semantic Activation architecture to simulate the N400 as the total activity produced within its semantic (output) layer (Laszlo & Plaut, 2012; Laszlo & Armstrong, 2014; Cheyette & Plaut, 2017). This model was able to simulate the effects of several lexical variables on the N400 produced by words presented in isolation, including the effects of orthographic neighborhood size (Laszlo & Plaut, 2012), lexical frequency (Cheyette & Plaut, 2017), and semantic richness (Cheyette & Plaut, 2017). It was also able to simulate the attenuation of the N400 to target inputs in both repetition (Laszlo & Armstrong, 2014) and semantic priming paradigms (Cheyette & Plaut, 2017), see Table 1.

**Table 1 -.** Phenomena simulated by computational models of the N400. An overview of the range of N400 phenomena that have been modeled in the literature.

|  |  | Word-level models |  |  |  | Sentence-level models |  |  | Predictive Coding |
| --- | --- | --- | --- | --- | --- | --- | --- | --- | --- |
|  |  | Laszlo & Plaut (2012) | Laszlo & Armstrong (2014) | Cheyette & Plaut (2017) | Rabovsky & McRae (2014) | Brouwer et al. (2017) | Rabovsky, Hansen & McClelland (2018) | Fitz & Chang (2019) |  |
| **Lexical variables** | Orthographic neighborhood size | ✓ | — | ✓ | ✓ | — | — | — | ✓ |
|  | Lexical frequency | — | — | ✓ | ✓ | — | ✓ | — | ✓ |
|  | Concreteness/Semantic richness | — | — | ✓ | ✓ | — | — | — | ✓ |
|  | Lexical status | — | — | — | — | — | — | — | ✓ |
| **Word-pair priming** | Repetition priming | — | ✓ | ✓ | ✓ | — | ✓ | ✓ | ✓ |
|  | Semantic priming | — | — | ✓ | ✓ | — | ✓ | — | ✓ |
|  | Associative priming | — | — | ✓ | — | — | ✓ | — | — |
| **Contextual effects** | Lexical probability | — | — | — | — | — | ✓ | ✓ | ✓ |
|  | Constraint for unexpected endings | — | — | — | — | — | ✓ | ✓ | ✓ |
|  | Anticipatory semantic overlap | — | — | — | — | — | ✓ | — | ✓ |
|  | Anticipatory orthographic overlap | — | — | — | — | — | — | — | ✓ |
|  | Role reversal anomaly | — | — | — | — | ✓ | ✓ | — | — |
|  | Semantic incongruity | — | — | — | — | ✓ | ✓ | — | — |
|  | Position in sentence | — | — | — | — | — | ✓ | ✓ | — |
|  | Word order violation | — | — | — | — | — | ✓ | — | — |
|  | Linguistic adaptation | — | — | — | — | — | ✓ | ✓ | — |
| **Interactions** | Constr. x Anticipatory sem. overlap | — | — | — | — | — | — | — | ✓ |
|  | Repetition x Frequency | — | — | ✓ | ✓ | — | — | — | ✓ |
|  | Repetition x Semantic Richness | — | — | ✓ | ✓ | — | — | — | ✓ |
|  | Repetition x Incongruity | — | — | — | — | — | ✓ | — | — |
|  | Cloze x Frequency | — | — | — | — | — | — | — | ✓ |
|  | Cloze x Semantic Richness | — | — | — | — | — | — | — | ✓ |

In another word-level model, Rabovsky and McRae (2014) simulated the N400 as the difference (cross-entropy error) between the activity produced within the model’s semantic (output) layer and an ideal “correct” semantic target presented to the model. They showed that this operationalization of the N400 could account for a similar range of findings as above (see Table 1).

The sentence-level models were trained to map a sequence of word inputs onto a higher event-level representation (Brouwer, Crocker, Venhuizen & Hoeks, 2017; Rabovsky, Hansen & McClelland, 2018), or onto the model’s prediction of a subsequent word (Fitz & Chang, 2019). These training goals required the model to retain a representation of the full sequence of prior inputs as well as to implicitly predict upcoming information. This was achieved by including a recurrent element in the network (cf. Elman & McClelland, 1984; Elman, 1990). The N400 was modeled either as “change in state” — the amount of change that the input induced within a particular hidden layer within the network (Brouwer, Crocker, Venhuizen & Hoeks, 2017; Rabovsky, Hansen & McClelland, 2018; Rabovsky, 2020), or as the difference between a next-word prediction that was explicitly generated by the model, and the word that was subsequently presented, termed “prediction error” (Fitz & Chang, 2019). Together, these models were able to simulate multiple effects of a prior context on the N400 evoked by incoming words (see Table 1, and Nour Eddine, Brothers & Kuperberg, 2022 for a detailed review).

The architectures and assumptions of these different computational models of the N400 are quite different from one another. However, it is worth emphasizing that in all except the Semantic Activation model (Laszlo & Plaut, 2012; Laszlo & Armstrong, 2014; Cheyette & Plaut, 2017), the N400 was operationalized as a *difference* that was computed by the modeler outside the model’s architecture. This difference value was conceptualized either as a prediction error (Rabovsky & McRae, 2014; Fitz & Chang, 2019, or as a change-in-state (Brouwer et al., 2017; Rabovsky et al., 2018) that emerged either as a byproduct of other computations (Brouwer et al., 2017) and/or served as a signal for downstream learning (Rabovsky et al., 2018; Rabovsky & McRae, 2014; Fitz & Chang, 2019). In no case, however, did it play a direct functional role in comprehension itself. This is in contrast with predictive coding, which proposes that prediction error, computed locally at each level of representation, plays an integral role in the optimization algorithm that the brain uses to approximate inference, i.e., the process of inferring meaning from an input’s linguistic form^1^.

Predictive coding was initially proposed to explain extra-classical receptive field effects in the visual cortex (Rao & Ballard, 1999; see also Mumford, 1992), and was later expanded into a more general account of perceptual inference in the brain (Friston, 2005; Clark, 2013; see also Spratling, 2016b). In predictive coding, *prediction error* is defined as the residual information observed at a given level of the cortical hierarchy that cannot be explained by top-down predictions (or “reconstructions’) that are generated by the level above. This error is encoded within “error units”, and is passed up to the level above where it is used to modify representations encoded within functionally distinct “state units”. As a result, these higher-level state units generate more accurate top-down predictions on the next iteration of the algorithm, which suppress the lower-level prediction error. This process takes place at each level of the hierarchy such that, over multiple iterations, the magnitude of prediction error –– the total activity produced by the error units –– gradually decreases as the state units converge upon the representation that best explains the bottom-up input.

The idea that higher levels of a representational hierarchy generate top-down predictions that facilitate the processing of inputs at lower levels is largely consistent with how prediction is typically framed in more general psycholinguistic and neurobiological frameworks of language processing. According to these accounts, during incremental language comprehension, the brain continually generates top-down predictions, based on a high-level interpretation of the prior context, which facilitate the processing of incoming words whose semantic features match these predictions (DeLong, Urbach, & Kutas, 2005; Federmeier, 2007; Kuperberg & Jaeger, 2016, section 3.5). The amplitude of the N400 can be conceptualized as reflecting the ease of accessing the semantic features of these incoming words (i.e., the ease of lexico-semantic access or retrieval), or as the amount of unpredicted lexico-semantic *information* encoded within the bottom-up input (see Kuperberg, 2016 for discussion). Indeed, some researchers have explicitly appealed to the principles of predictive coding to explain the functional role of the N400 (e.g. Xiang & Kuperberg, 2015; Rabovsky & McRae, 2014; Bornkessel-Schlesewsky & Schlesewsky, 2019; Kuperberg, Brothers, & Wlotko, 2020). To date, however, there have been no attempts to simulate the N400 (or any other language ERP component) using an implemented computational model of predictive coding.

In the present work, we built a computational model of lexico-semantic processing that was based on the same predictive coding principles as those used to simulate low-level neural phenomena in the visual system (Rao & Ballard, 1999; Spratling, 2012; Spratling, 2013; Spratling, 2014). We operationalized the N400 as *lexico-semantic prediction error ––* the total activity produced by error units at the semantic and lexical levels on each iteration of the algorithm, as the model inferred the meaning of orthographic inputs. We carried out a series of simulations to determine whether the principles of predictive coding can account for the temporal dynamics of the N400, as well as its functional sensitivity to (1) various lexical variables, (2) priming, (3) contextual effects, and (4) their higher-order interactions.

## Methods

### Model Architecture

The basic structure of the hierarchical predictive coding model used in all simulations is shown in Figure 1. It consisted of four layers — three levels of linguistic representation (orthographic, lexical, semantic) and a fourth “dummy” layer at the top, which was used to simulate the top-down effects of context.

**Figure 1 -.**
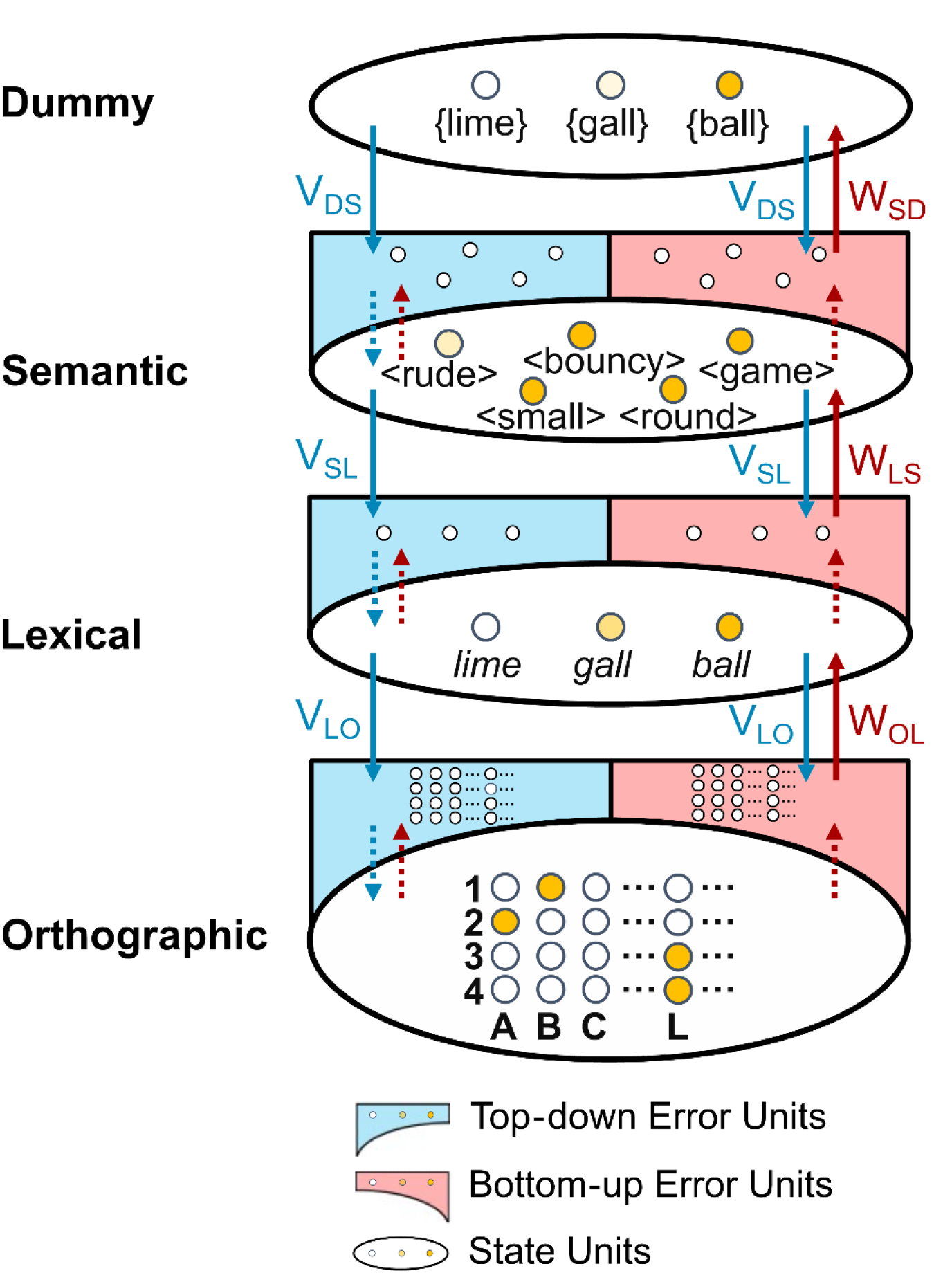
Predictive coding model architecture. State units at three levels of linguistic representation (Orthographic, Lexical and Semantic) and at the highest dummy layer are depicted as small circles within the large ovals. Error units at each of the three levels of linguistic representation are depicted as small circles within the half arcs. Dotted arrows indicate one-to-one connections between error and state units at the same level of representation. Solid arrows indicate many-to-many connections between error and state units across levels of representation. These many-to-many connections were specified using hand-coded weight matrices: W (feedforward) and V (feedback). V_LO_/W_OL_: Connections between the lexical and orthographic level; V_SL_/W_LS_: Connections between the semantic and lexical level; V_DS_/W_SD_: Connections between dummy and semantic level. We schematically depict the activity pattern of the model’s state units after it has settled on the representation of the item, “ball”. Different shades of yellow are used to indicate each state unit’s strength of activity. At the Orthographic level, four state units are activated: B in the first position, A in the second position, and L in the final two positions. At the Lexical level, the unit corresponding to *ball* is mostly strongly activated, and its orthographic neighbor *gall* is partly activated because it shares three letters with *ball*. At the Semantic level, the units corresponding to the semantic features of *ball* (<bouncy>, etc.) are shown with different levels of activation. At the highest Dummy layer, the unit corresponding to the representation of “ball” is most strongly activated. Because the model has settled, activity within error units at all levels is minimal.

Consistent with predictive coding principles, each of the three levels of linguistic representation had two populations of connectionist units — *state units*, which encode the internal representations being inferred by the predictive coding algorithm, and *error units,* which encode the *difference* in information (i.e., residual information) between that represented by state units at this level and top-down predictions (otherwise referred to as “reconstructions”) generated by state units at the level above. We incorporated two types of error units (Spratling, 2016b): (a) “bottom-up error units”, which represent the residual information encoded in the state units that was not encoded in the top-down reconstructions (known as “prediction error”), and (b) “top-down error units”, which represent the residual information encoded in the top-down reconstructions that was not encoded in the state units (termed “top-down bias”). As discussed further below, prediction error and top-down bias each play a different role in the predictive coding algorithm.

The lowest *orthographic* level included 104 sets of state/error units, each encoding one of 26 letter identities (A-Z) at one of four possible spatial positions (following McClelland & Rumelhart, 1981). The middle *lexical* level included 1,579 sets of state/error units, each representing a four-letter word in the model’s lexicon (e.g., *baby, lime*). 512 of these lexical units corresponded to the 512 *critical words* that we used for most of our simulations. By design, the orthographic neighborhood size, frequency and semantic richness (operationalized as described below) of these critical words were uncorrelated. The third *semantic* level included 12,929 state/error units, each representing a unique semantic feature (e.g., <small>, <human>; following Cheyette & Plaut, 2017; Rabovsky & McRae, 2014). Finally, the highest dummy layer of the hierarchy, which only had state units, collectively represented a probability-like distribution over the lexicon, allowing the modeler to provide the model with pre-activation to simulate the top-down effects of context.

The connections within and across layers of the model were prescribed by predictive coding principles. *Within* each linguistic level, the state and error units shared one-to-one connections. *Across* successive levels, each state unit was connected to error units via many-to-many feedback/feedforward connections. In other words, across levels, state units communicated exclusively through error units.

The many-to-many connections that link state units at a higher level with error units at a lower level were hand-coded using two matrices, V and W. The *feedback* connections were specified by matrix V, which specified the *generative parameters* of the model; that is, each column of the V matrix specified how a given higher-level “latent cause” generated an idealized noise-free pattern of observations at the lower level. The *feedforward* connections were specified by matrix W, which was simply the transpose of matrix V.

The *orthographic-lexical* matrices connected lower-level orthographic error units with higher-level lexical state units. These matrices specified the *spelling* of each word; that is, each column in the V matrix specified the mapping between a particular lexical item and the correct position of each letter (e.g., *ball →* B in position 1, A in position 2, L in position 3, and L in position 4; see Figure 2). This resulted in each lexical item having a particular orthographic neighborhood size. For the purpose of our simulations, we defined the orthographic neighborhood size of each item as the number of lexical units with which it shared 3 letters. For example, *ball* and *gall* are orthographic neighbors because they share A_2_, L_3_, and L_4_, see Figure 2.

**Figure 2 -.**
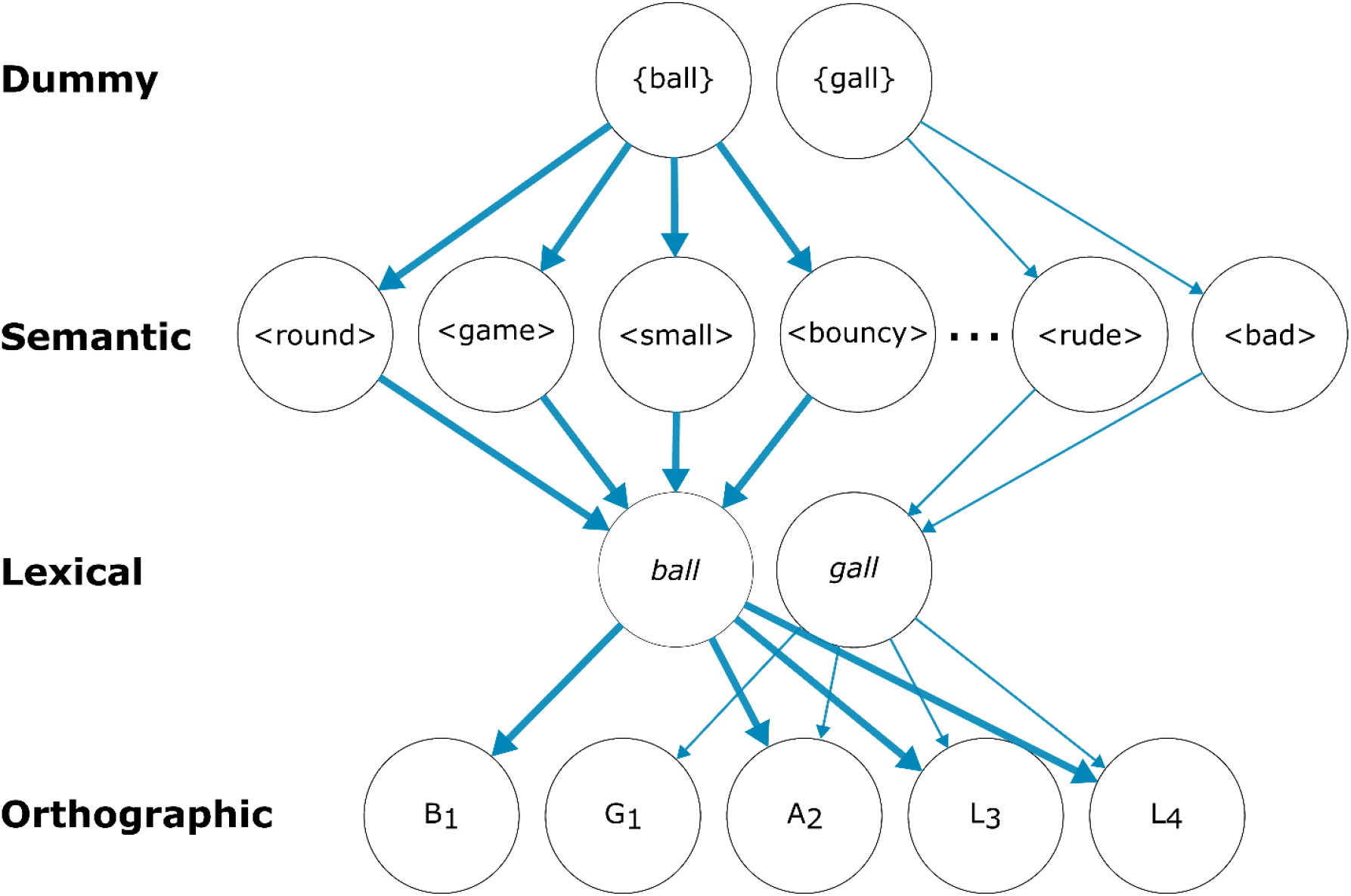
Schematic illustration of the feedback connections for two words in the model’s lexicon. Each circle indicates a representational node, and the blue arrows indicate feedback connections between layers. Note that, for simplification, this diagram does not distinguish between state and error units. In the model itself, however, the feedback connections linked higher-level state units with lower-level error units. To specify the frequency of each lexical item, we modified the connection strengths of its unique set of feedback connections. This is depicted schematically using arrow thickness. For example, the arrows are thicker for *ball* than *gall* because *ball* is more frequent. Although each lexical item has its own unique set of connections, these connections can terminate on shared nodes. For example, the lexical-orthographic feedback connections for *ball* and *gall* both terminate on the same A_2_, L_3_, and L_4_ nodes, and the semantic-lexical feedback connections for the semantic features, <round>, <game>, <small> and <bouncy>, all terminate on the same lexical node, *ball*. In the model itself, this resulted in each lexical item having a particular “orthographic neighborhood size” and a particular “semantic richness”. For example, *ball* and *gall* are orthographic neighbors, and the semantic richness of the word *ball* is greater than *gall* because the former lexical item is connected to more semantic features (4 *vs*. 2). For the purpose of our simulations, we defined each lexical item’s orthographic neighborhood size as the number of lexical units with which it shared 3 letters. We defined “semantically rich” items as those that were connected to 18 features, and “non-rich” items as those that were connected to 9 features.

The *lexico-semantic* matrices connected lower-level lexical error units with higher-level semantic state units. These matrices specified the *meaning* of each word; that is, each row in the V matrix specified the mappings between each lexical unit and its particular set of semantic features (e.g., *ball* → <round>, <bouncy>, etc.). To define the semantic richness of each of the 512 critical lexical units used in most of the simulations, we assigned half of these items 18 semantic features (semantically *rich* items) and the other half 9 semantic features (*non-rich* items), based on a median split on their concreteness ratings taken from Brysbaert, Warriner, and Kuperman (2014). Each of these 512 words shared between 0 and 8 semantic features with at least one other lexical unit, allowing us to simulate effects of semantic relatedness on the N400. Each of the remaining 1,067 lexical units was assigned 9 unique semantic features.

In order to set the frequency of lexical items within the model, we modified the strength of each item’s unique set of feedback connections (see Figure 2). This is because lexical frequency information is acquired gradually over time, and is thought to bias perceptual inference by providing a top-down prior (Norris, 2006). Specifically, we increased the value of each word’s entries in the V matrices with a value that was proportional to its SUBTLEX-US frequency (Brysbaert & New, 2009)). This score was obtained by shifting and scaling the log frequencies of all 1,579 words in the model lexicon so that they fell into the [0, 0.1] range. This range was selected so that the addition of frequency scores did not dramatically alter the mean and maximum value of the connections in any matrix.

### The Predictive Coding Algorithm

Predictive coding is an optimization algorithm for approximating Bayesian inference. In the current setting, this involves finding the combination of latent causes (i.e., columns of V matrices) that yields the actual observed pattern (see Spratling, 2017; Spratling, 2016a). Here we implemented a modified version of the Predictive Coding/Biased Competition-Divisive Input Modulation algorithm (Spratling, 2008; Spratling, 2016b). This algorithm shares many processing principles with the influential predictive coding approach developed by Rao & Ballard (1999). However, the error units compute the residual information via element-wise division rather than element-wise subtraction (see Spratling, 2008). This ensures rapid convergence of the algorithm and guarantees that the activity across all units remains non-negative, similar to biological neurons.

At each iteration, *n,* of the predictive coding algorithm, the following processes occur in sequence at each level of the hierarchy. This is illustrated schematically in Figure 3.

**Figure 3 -.**
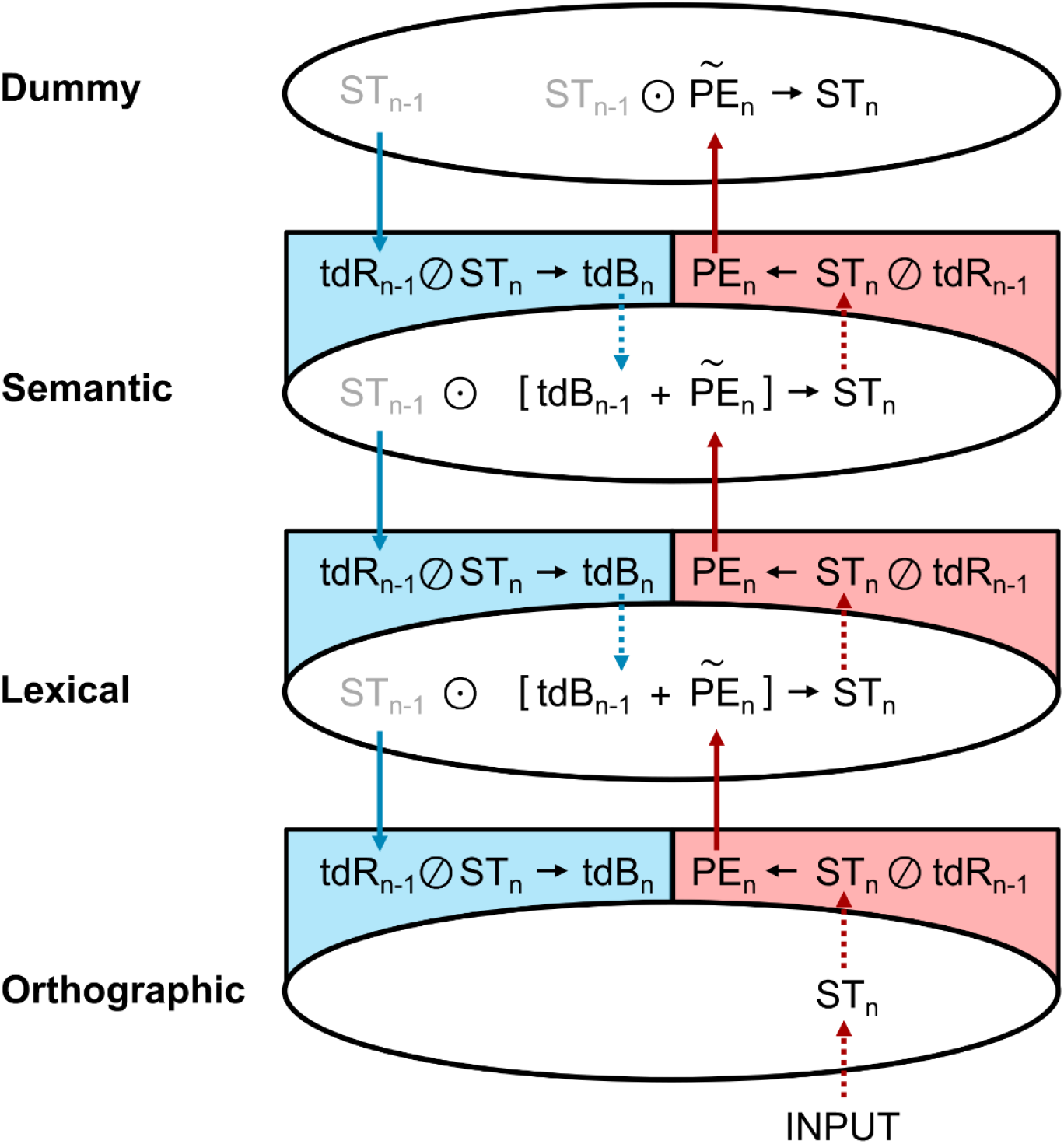
Predictive coding algorithm. Schematic illustration of the predictive coding algorithm operating on the *n*^th^ iteration, following the presentation of bottom-up orthographic input. As in Figure 1, at each layer, the large ovals contain state units, the red half-arcs contain bottom-up error units, and the blue half-arcs contain top-down error units. Each variable’s subscript indicates the iteration on which it was computed. Solid arrows indicate the linear transformation of a variable through the V and W matrices. Dotted arrows indicate the copying of a variable. The same three steps occur in sequence at each level of representation: (1) State units are updated, based on (a) the top-down bias computed at the same level on the previous iteration, and (b) the prediction error computed at the level below on the same iteration 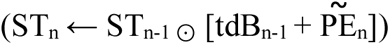, and their values are copied to the top-down and bottom-up error units at the same level. (2) Bottom-up error units compute prediction error (PE_n_) through elementwise division (ST_n_ Ø tdR_n-1_) and pass this prediction error up to state units at the level above by transforming its dimensionality 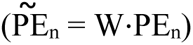, and top-down error units compute top-down bias (tdB_n_) and copy this top-down bias to state units at the same level so that it is ready to update the state units on the subsequent [*n*+1]^th^ iteration); (3) State units generate top-down reconstructions of activity at the level below via linear transformation by the V (generative) matrix, i.e., V⋅ST = tdR, and pass these reconstructions down to the error units at the level below.

#### The updating of state units

The state units at each level represent the information that is being inferred, regardless of its predictability. At each level, state units are updated based on (a) the bottom-up input on the current iteration (*n*) and (b) a top-down bias (tdB) that was computed on the previous iteration (*n*-1). At the orthographic level, the bottom-up input is the orthographic vector provided by the modeler; at higher levels, the input is the prediction error (PE) that is computed at the level below on the current iteration of the algorithm. The updated state is computed through element-wise multiplication: 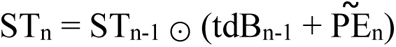.

Note that prediction error and top-down bias each play different roles in this state update. Prediction error, which is computed at the level below on the same iteration of the algorithm, allows these states to be updated so that they will generate more accurate top-down reconstructions of activity at the level below on the subsequent iteration of the algorithm. In contrast, the top-down bias, which was computed at the same level on the previous iteration of the algorithm, modifies the state pattern such that it is brought closer to the top-down reconstruction that was generated by the level above on the previous iteration, i.e., so that this state pattern serves as a better “target” for these higher-level reconstructions/predictions on the current iteration of the algorithm.

#### The computation of residual information: Prediction error and Top-down bias

The error units at each level represent the residual *difference* in information between the top-down reconstruction that was computed on the previous iteration of the algorithm (tdR_n-1_) and the updated state pattern at the same level on the current iteration (ST_n_). As noted earlier, we included two types of error units — bottom-up error units and top-down error units.

Within the bottom-up error units, the updated state pattern (at the same level) is divided elementwise by the top-down reconstruction (tdR) from the level above (which was computed on the previous iteration of the algorithm), yielding prediction error: PE_n_ = ST_n_ Ø tdR_n-1_. This prediction error encodes the residual information within the state units that is not present within the top-down reconstruction. It is multiplied by the W matrix 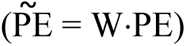, which transforms its dimensionality so that it can serve as the bottom-up input for updating state units at the level above on the current iteration of the algorithm.

Within the top-down error units, the top-down reconstruction, which was computed on the previous iteration of the algorithm, is divided elementwise by the current state through element-wise division, yielding a top-down Bias (tdB): tdB_n_ = tdR_n-1_ Ø ST_n_. This top-down bias encodes residual information within the top-down reconstructions that is not present within the state units. It is passed to state units at the same level through one-to-one connections, and is used to bias state updates on the subsequent iteration of the algorithm (*n*+1).

#### The generation of top-down reconstructions (predictions) to be used on the subsequent iteration

After the state units are updated at all levels, they generate a top-down reconstruction (tdR) of the pattern of activity at the level below by multiplying the current higher-level state vector (ST_n_) by the V (generative) matrix, i.e., V⋅ST_n_ = tdR_n_. This top-down reconstruction will be used to compute the prediction error and the top-down bias on the subsequent iteration of the algorithm (as described under step 2).

Thus, over multiple iterations of the algorithm, as higher-level states are updated, both the magnitude of the prediction error and the top-down bias decrease, and the model reaches a global, internally consistent state that can accurately explain the bottom-up input at its multiple levels.

### Simulations and visualizations

We simulated a wide range of benchmark phenomena in the N400 literature. These included: (1) Lexical effects (effects of orthographic neighborhood size on words and pseudowords, and effects of lexical frequency and semantic richness on words); (2) Priming effects (repetition priming and semantic priming); (3) Contextual effects (lexical probability, contextual constraint, effects of anticipatory semantic overlap on words, and effects of anticipatory orthographic overlap on words and pseudowords), and (4) Interactions of each of the lexical variables with repetition priming and lexical probability.

For simulations involving only real words, we used orthographic input vectors that corresponded to 512 four-letter “critical words” in the model’s lexicon. These words were selected to have a wide range of lexical frequencies, orthographic neighborhood sizes, and semantic richness values (half rich, half non-rich), and, by design, these three variables were all uncorrelated (|*r*| < 0.07) across our 512 critical words. For simulations involving pseudowords, we used 400 words and 400 pseudowords that were matched on orthographic neighborhood size (for details of how this stimulus set was constructed, see Results, section 3.6).

Details of how we carried out each specific simulation are given in the Results section. In all simulations, after initializing the model, we presented the bottom-up orthographic input by clamping an orthographic vector that corresponded to the stimulus of interest. We then ran the predictive coding algorithm for 20 iterations. On each iteration following stimulus onset, we computed the total activity produced by the lexical and semantic error units, i.e., *lexico-semantic prediction error*, and averaged these values across items. In this way, we constructed a full time course of the simulated N400.

### Statistical analyses

For all statistical analyses, we operationalized the amplitude of the simulated N400 as the mean magnitude of the total lexico-semantic prediction error produced by each item, averaged across iterations 2 to 11 following stimulus presentation. In all cases, this constituted a 10-iteration time window that surrounded the peak of the error response.

To examine the effects of lexical variables, which varied between items, we carried out simple regression analyses. To examine the effects of priming and contextual variables, we carried out linear mixed effects regressions (lmer), using lme4 package version 1.1-31(Bates et al., 2015) in R version 4.2.2 (R Core Team, 2022). In these models, we first attempted to fit the maximal random effects structure. In the case of convergence failures, we simplified the random effects structure following the recommendations of Barr et al. (2013). Statistical significance was assessed using a type-III sums of squares estimation, with p-values estimated using the Satterthwaite approximation (Satterthwaite, 1946) using lmerTest version 3.1-3 (Kuznetsova et al., 2017).

## Results

### Time course of the simulated N400

In all simulations, the time course of the simulated N400 (lexico-semantic prediction error) showed a rise-and-fall waveform-like morphology, similar to the empirical N400. After stimulus onset, the magnitude of the lexico-semantic prediction error rose to a peak at around iteration 5 before steadily decreasing to a minimum by iteration 20.

### 1. Effects of lexical variables

#### 1.1 Effect of orthographic neighborhood size on words

Empirically, words with more orthographic neighbors (e.g. *ball*: *bull*, *call*, *bail*) produce a larger N400 response than words with fewer neighbors (e.g., *kiwi*; Holcomb, Grainger, & O’Rourke, 2002; Laszlo & Federmeier, 2011).

In our model, orthographic neighborhoods were determined by the pattern of weights (specified by the V and W matrices) that connected the lexical and orthographic units (see Figure 2 in Methods). For our simulations, we operationalized the *orthographic neighborhood size* (ONsize) of each of our 512 critical lexical items as the number of words in the model’s lexicon with which it overlapped in three letter positions. Mirroring the empirical findings, we found that lexico-semantic prediction error was larger on words with a larger *versus* smaller ONsize (*b* = 32.63, *t* = 43.82, *p* < .001, see Figure 4A).

**Figure 4 -.**
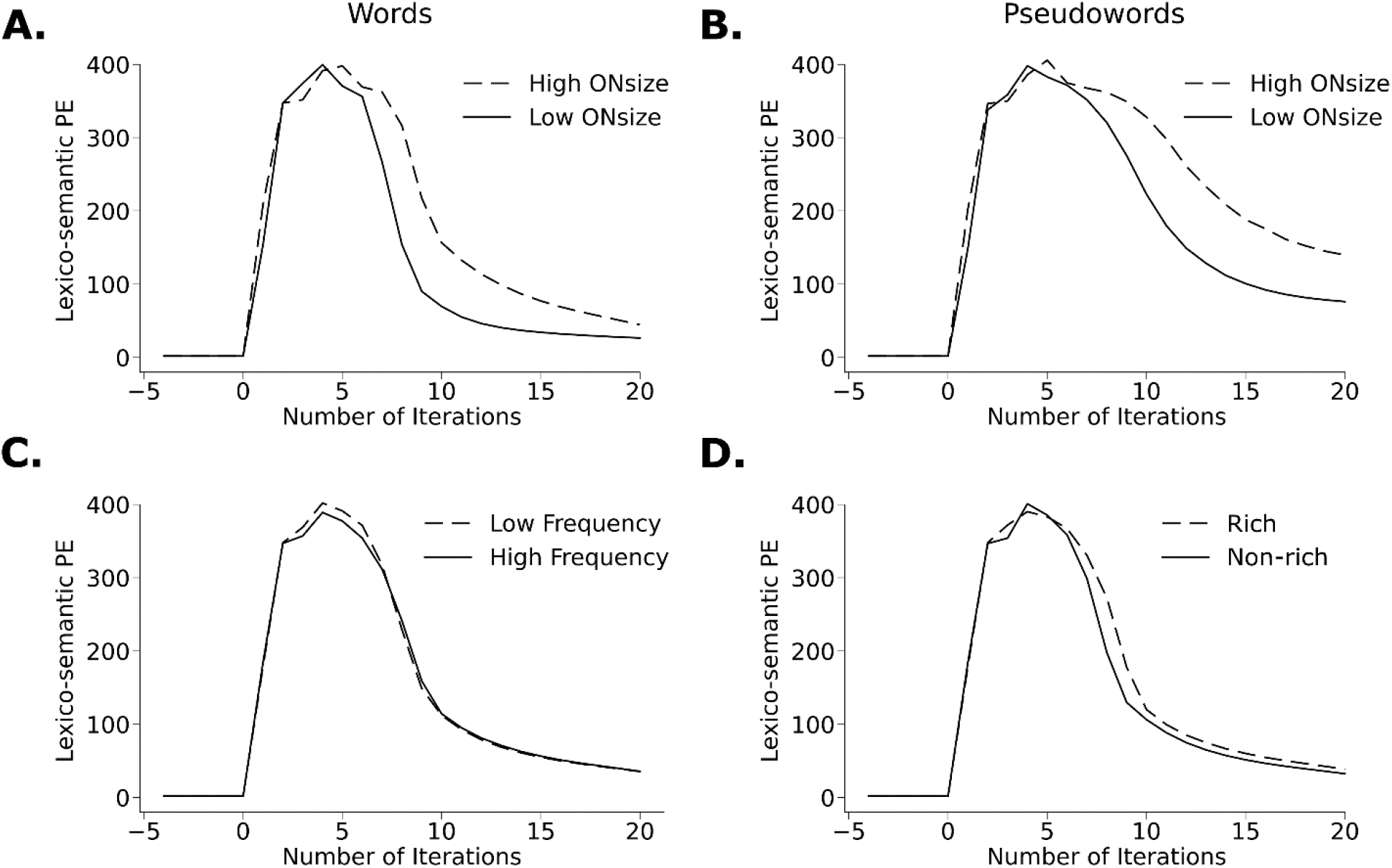
Effects of lexical variables on the time course of lexico-semantic prediction error. On each plot, the x-axis shows the number of iterations since stimulus onset, and the y-axis shows the total lexico-semantic prediction error (PE) (arbitrary units), averaged across items within each condition. **A.** High *vs.* Low Orthographic Neighborhood size (ONsize), based on a median split across 512 critical words. High ONsize words elicited a significantly larger lexico-semantic prediction error than Low ONsize words. **B.** High *vs.* Low ONsize, based on a median split across 400 pseudoword items. High ONsize pseudowords elicited a significantly larger lexico-semantic prediction error than Low ONsize pseudowords. **C.** High *vs.* Low Frequency, based on a median split across 512 critical words. Low frequency items elicited a significantly larger lexico-semantic prediction error than high frequency items. **D.** Rich *vs.* Non-rich (lexical items connected to 18 *vs.* 9 semantic features). Rich items elicited a significantly larger lexico-semantic prediction error than Non-rich items.

#### 1.2 Effect of orthographic neighborhood size on pseudowords and effect of lexical status

Empirical studies have shown that orthographic neighborhood size not only modulates the amplitude of the N400 produced by real words (WISH), but also by pseudowords (*WUSH, Laszlo & Federmeier, 2011; Holcomb, Grainger & O’Rourke, 2002). Moreover, the magnitude of this effect is the same on words and pseudowords (Laszlo & Federmeier, 2011). On the other hand, several studies have reported that pseudowords elicit larger N400s than words (“bave” > “gave”; Bentin, 1987), even when controlling for orthographic neighborhood size (Holcomb, Grainger & O’Rourke, 2002; Meade, Midgley, Dijkstra & Holcomb, 2018; Braun, Jacobs, Hahne, Ricker, Hofmann & Hutzler, 2006; although see Laszlo & Federmeier, 2011).

To determine whether our model could explain these effects of orthographic neighborhood size on the N400 produced by pseudowords, and to examine the effect of lexical status (words *vs.* pseudowords) on lexico-semantic prediction error, we carried out another set of simulations, this time using a new set of 400 words and 400 pseudowords that were designed to have identical neighborhood sizes (see section 3.6 below for details of how these stimuli were developed). Consistent with the empirical findings, we saw an effect of orthographic neighborhood size on pseudowords (see Figure 4B). Consistent with the empirical findings, this effect did not differ between the words and pseudowords (Main effect of ONsize: *b =* 0.50, *t =* 27.03, *p* < .001; no interaction between ONsize and Lexical Status: *b* = -0.002, *t =* -0.09, *p =* .93). Also consistent with many of the empirical findings (Holcomb, Grainger & O’Rourke, 2002; Meade, Midgley, Dijkstra & Holcomb, 2018; Braun et al., 2006), there was also a main effect of Lexical Status due to a larger lexico-semantic prediction error on pseudowords than words (*b =* -0.69*, t =* -36.77*, p <* .001).

These effects of orthographic neighborhood size on both words and pseudowords arose because when the bottom-up input arrived at the lexical level, it partially activated a large number of closely overlapping lexical state units, which led to the increased propagation of prediction error to the lexical and semantic levels of representation. The reason why pseudowords (WUSH) produced a larger lexico-semantic prediction error than words, even when controlling for orthographic neighborhood size, is that, after activating multiple lexical candidates with varying degrees of orthographic overlap (WISH, BUSH, LUSH, etc.), the model was unable to settle on a single lexico-semantic state that could explain the bottom-up input.

#### 1.3 Effect of lexical frequency

The amplitude of the N400 is smaller in response to words of higher frequency (e.g., “ball”) than lower frequency (e.g. “gall”; Rugg, 1990; Van Petten & Kutas, 1990; Laszlo & Federmeier, 2014; Hauk et al., 2006). As described under Model Architecture, and depicted schematically in Figure 2, we incorporated the frequency of each word in the model by increasing the strength of its unique set of feedback connections in proportion to its SUBTLEX-US frequency (Brysbaert & New, 2009)). This ensured that units associated with higher frequency items received stronger feedback from higher-level state units, providing a top-down prior during perceptual inference (see Norris, 2006).

As expected, our simulations showed that higher frequency words produced a smaller lexico-semantic prediction error than lower frequency words (*b* = -4.05, *t* = -5.43, *p* < .001; see Figure 4C). This is because the stronger feedback weights allowed higher levels of the network to generate predictions that better suppressed the production of lexical and semantic prediction error.

#### 1.4 Effect of semantic richness

The N400 is generally larger to words with more concrete meanings (Kounios & Holcomb, 1994; Holcomb, Kounios, Anderson & West, 1999; Lee & Federmeier, 2008), more semantic associates (Laszlo & Federmeier, 2011), and a larger number of semantic features (Amsel, 2011; Rabovsky, Sommer & Abdel Rahman, 2012b; but see Kounios, Green, Payne, Fleck, Grondin & McRae, 2009).

Following previous models (Cheyette & Plaut, 2017; Rabovsky & McRae, 2014), to capture these effects, we operationalized the semantic richness of each word in the model’s lexicon as the number of semantic features with which it was connected. As described under Model Architecture (see Methods), each of our 512 critical lexical items was connected to either 18 features (Rich) or 9 features (Non-rich).

Our simulations show that semantically rich words produced a larger lexico-semantic prediction error than the Non-rich words (Richness: *b* = 9.03, *t* = 12.12, *p* < .001; see Figure 4D). This follows from the simple fact that prediction error is summed elementwise across all error units. Therefore, if a lexical unit is linked to a larger number of semantic units, its activation will produce a larger total prediction error^2^.

### 2. Effects of word-pair priming

In a typical priming paradigm, pairs of “prime” and “target” words are presented sequentially, with the prime being either related or unrelated to the target along some dimension. Empirically, the amplitude of the N400 is smaller to repeated than non-repeated targets in repetition priming paradigms (e.g. *lime* – *lime* vs. *flow* – *lime*: Rugg, 1985; Misra & Holcomb, 2003) and smaller to semantically related than unrelated targets in semantic priming paradigms (e.g. *sour – lime* vs. *flow – lime:* Bentin, McCarthy &Wood, 1985; Rugg, 1985; Holcomb, 1988; Holcomb & Neville, 1990). Additionally, several studies have shown that the repetition priming effect is larger than the semantic priming effect (Rugg, 1985; Deacon, Dynowska, Ritter & Grose-Fifer, 2004).

To simulate these priming effects, we clamped an orthographic “prime” input for 20 iterations, followed by two blank iterations (all input units clamped to zero), followed by either a related or unrelated “target” input for an additional 20 iterations.

#### 2.1 Effect of repetition priming

In the repetition priming simulations, the prime was either identical or unrelated to the target (sharing no semantic features). Consistent with the empirical findings, we found that the repeated targets produced a smaller lexico-semantic prediction error than the non-repeated targets (*b* = -125.52, *t* = -140.3, *p* < .001; see Figure 5A). This is because the presentation of the prime word led the model to fully converge on the lexical and semantic state units that corresponded to the target. Therefore, when the repeated target was presented, the lexico-semantic predictions were largely accurate, and so very little error was produced.

**Figure 5 -.**
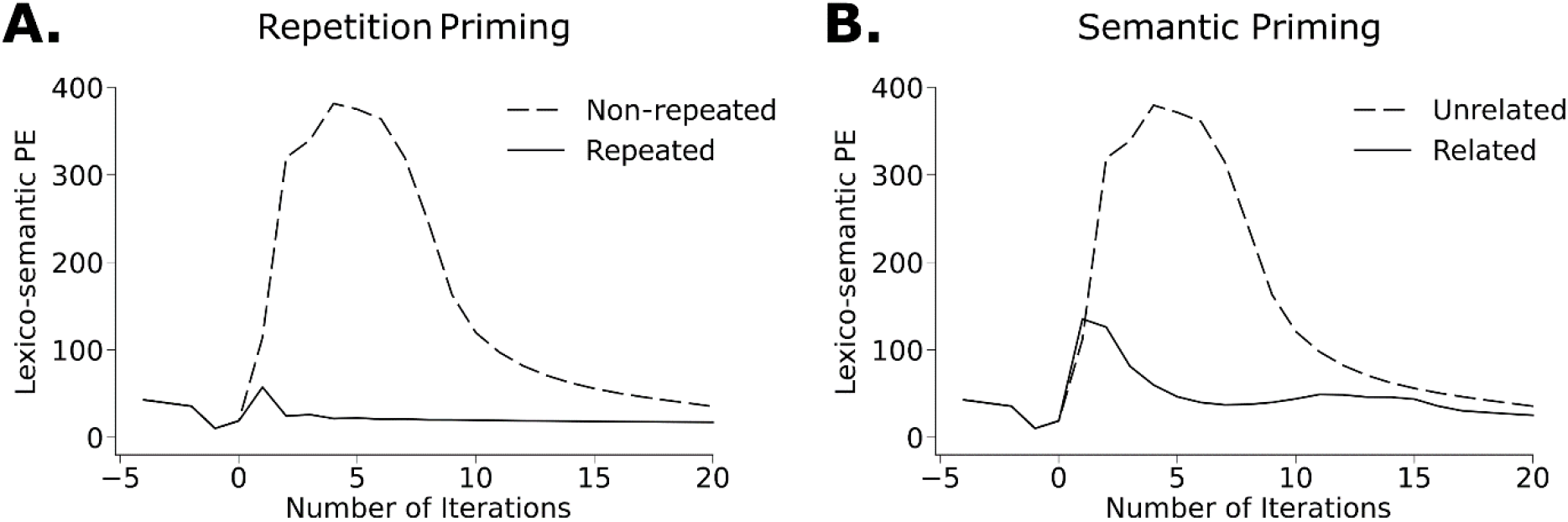
Effects of word-pair priming on the time course of lexico-semantic prediction error. A. Effect of repetition priming. **B.** Effect of semantic priming: Unrelated (zero semantic features shared between prime and target) *vs.* Related (eight semantic features shared between prime and target).

#### 2.2 Effect of semantic priming

In the semantic priming simulations, the prime either shared 8 semantic features (related condition) or no semantic features with the target (unrelated condition). Again consistent with the empirical findings, we found that the semantically related targets produced a smaller lexico-semantic prediction error than the unrelated targets (*b* = -107.25, *t* = -87.15, *p* < .001; see Figure 5B). This is because the presentation of the semantically related prime led the model to partially converge on the lexical and semantic state units that corresponded to the target, resulting in more accurate top-down predictions and therefore a smaller lexico-semantic prediction error to related targets.

Finally, consistent with the empirical findings, lexico-semantic prediction error was smaller to repeated than to semantically related targets (*b* = -34.67, *t* = -20.46, *p* < .001).

### 3. Contextual effects

During language comprehension, the amplitude of the N400 is strongly influenced by the broader sentence and discourse context. Predictive models of online language comprehension posit that this is because comprehenders incrementally use the prior context to infer a higher-level interpretation, which they use to generate top-down predictions that *pre-activate* anticipated lexico-semantic representations (e.g., Federmeier, 2007; Kuperberg & Jaeger 2016, section 3.5; Wang, Kuperberg & Jensen, 2018).

We carried out several simulations to model various top-down contextual effects on the N400. In all these simulations, we provided the model with top-down pre-activation by clamping the expected state unit at the highest dummy layer of the model with the desired percentage of pre-activation. We did this by fixing the total level of activation to a constant value (see Supplementary Materials) and distributing the remaining activation evenly across the remaining units. During this pre-activation phase, the steps of the predictive coding algorithm remained unchanged: the dummy layer generated semantic reconstructions, which induced a top-down bias in the model’s semantic state units. These newly activated semantic states then produced reconstructions that led to the pre-activation of lexical state units, and so on down the network. After 20 iterations of pre-activation, we unclamped the expected dummy state units at the top of the network and presented a new bottom-up input at the orthographic level for an additional 20 iterations.

#### 3.1 Effects of lexical probability

The amplitude of the N400 is strongly influenced by the lexical *probability* of each word, given its prior context. The probability of a word can be estimated either using the cloze procedure (the proportion of participants who produce that word during a sentence completion task (Taylor, 1953), or using large language models (e.g., GPT-3, Brown et al., 2020). Several studies have shown that the amplitude of the N400 is inversely proportional to these estimates of lexical probability in context (studies using cloze: Kutas & Hillyard, 1984; DeLong, Urbach & Kutas, 2005; Wlotko & Federmeier, 2012; studies using large language models: Michaelov, Coulson, & Bergen, 2021; Szewczyk & Federmeier, 2022; Heilbron, Armeni, Schoffelen, Hagoort & De Lange, 2022).

To simulate the effect of lexical probability on the N400, we presented each of our 512 critical words at four different levels of probability: 99%, 50%, 25% and uniform (1/[total words] = 1/1579 = 0.06%). In the lmer analyses, Probability was standardized and served as a within-item predictor. As expected, we observed a graded reduction in lexico-semantic prediction error as lexical probability increased (*b* = -87.42, *t* = -54.02, *p* < .001; see Figure 6A). This is because, with increasing predictability, the model was able to settle on an increasingly more accurate set of semantic and lexical states prior to the appearance of the target word. After stimulus onset, this resulted in more accurate top-down predictions and greater suppression of lexico-semantic prediction error.

**Figure 6 -.**
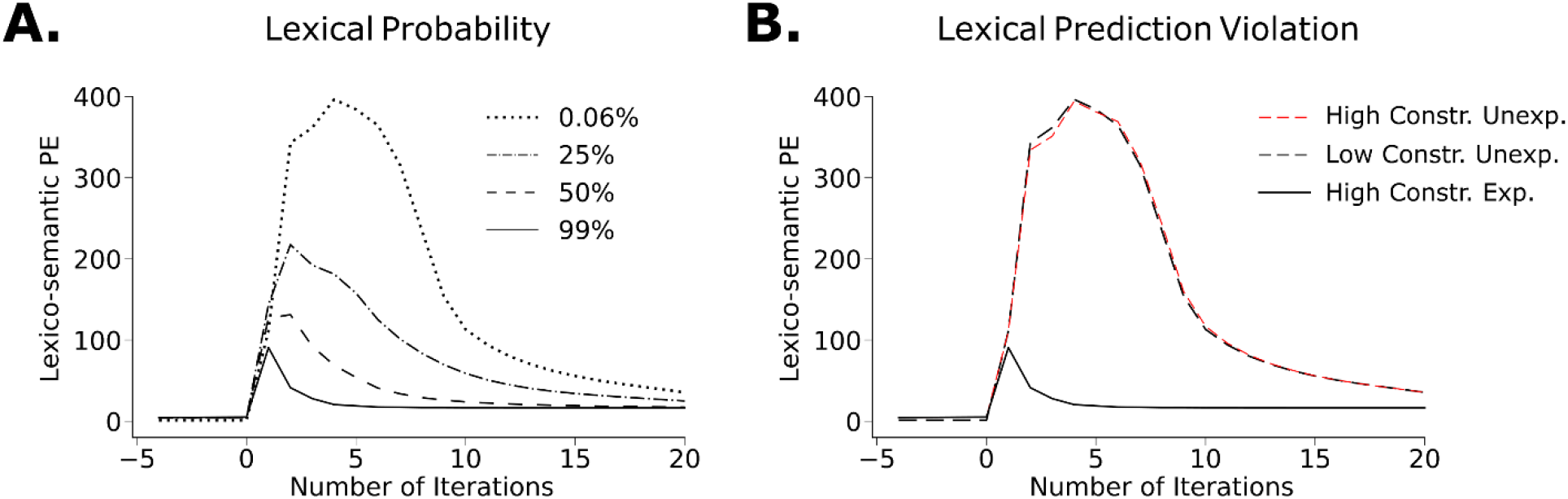
Effects of Lexical Probability and Constraint on the time course of lexico-semantic prediction error. A. Effect of lexical probability: Lexico-semantic prediction error decreased with increasing lexical probability. **B.** Effect of Constraint. Lexico-semantic prediction error was equally large to *high constraint unexpected* (HighConstr. Unexp.) and *low constraint unexpected* (Low Constr. Unexp.) inputs, relative to the *expected* inputs (High Constr. Exp).

#### 3.2 No effect of Constraint on unpredicted words

The amplitude of the N400 is not sensitive to the lexical constraint of the prior context, when controlling for lexical probability. For example, the N400 response is equally large to unexpected but plausible words that violate strong lexical predictions in a high constraint contexts (e.g. “*Every morning he took his dog for a <u>swim</u>*”, where the word “*walk*” was expected) and to unexpected but plausible words in low constraint contexts (e.g. “*Helen reached up to dust the <u>dresser</u>*”) (Kutas & Hillyard, 1984; Federmeier, Wlotko, De Ochoa-Dewald & Kutas, 2007; Kuperberg, Brothers, & Wlotko, 2020).

We simulated this finding by presenting an unexpected critical word in either a *high constraint* condition, in which we strongly pre-activated a different randomly selected word (99%), or in a *low constraint* condition in which all words were given uniform pre-activation (0.06%). In an lmer analysis with Constraint as a categorical within-items predictor, we found that there was no difference in the magnitude of lexico-semantic prediction error produced by the *high constraint unexpected* and the *low constraint unexpected* inputs (Constraint: *b* = 0.23, *t* = 0.85, *p* = .40; see Figure 6B). This is because lexico-semantic prediction error is only sensitive to the residual lexico-semantic information encoded within the bottom-up input that was *not* predicted by the level above. The amount of residual lexico-semantic information encoded within an unexpected input is the same, regardless of whether prior incorrect predictions were concentrated over one specific set of semantic features/lexical candidate, or whether they were spread diffusely over multiple semantic features/lexical candidates.

#### 3.3 Effect of anticipatory semantic overlap

In addition to its sensitivity to lexical probability, the amplitude of the N400 is also sensitive to the semantic relationship between a predicted word and the observed bottom-up input (Kutas & Hillyard, 1984; Federmeier & Kutas, 1999). For example, Federmeier and Kutas (1999) presented participants with highly constraining contexts, e.g. “*They wanted to make the hotel look more like a tropical resort. So along the driveway, they planted rows of*…”, followed by critical words that were highly predictable (e.g., *palms*), lexically unpredictable (<1% probability) but with semantic features that overlapped with the expected continuation (e.g. *pines*), or lexically unpredictable (<1% probability) but with fewer overlapping expected features (e.g. *tulips*). They observed a graded reduction in the N400 response across the three conditions (*palms* < *pines* < *tulips*).

To simulate this *anticipatory semantic overlap effect*, we pre-activated the model with each of our 512 words, assigning the dummy states a probability of 99% and clamping them for 20 iterations. We then presented the model with (a) the same word that was pre-activated (*expected*), (b) a different word that shared eight semantic features with the expected word (*unexpected semantically overlapping*), or (c) a different word that shared no semantic features with the expected word (*unexpected unrelated)*. We also ensured that the *unexpected semantically overlapping* and the *unexpected unrelated* words had minimal orthographic overlap with the *expected* words (0.38 and 0.37 characters respectively) and that the extent of this overlap did not differ between the two conditions (*t* < 1, *p* = .73).

As shown in Figure 7A, consistent with the empirical findings, we saw a graded reduction of lexico-semantic prediction error across the three conditions, with the *unexpected unrelated* words producing a significantly larger prediction error than the *unexpected semantically overlapping* words (*b =* 158.07; *t* = 61.42, *p* < .001), which, in turn, produced a significantly larger prediction error than the *expected* words (*b =* 95.27; *t* = 37.02, *p* < .001).

**Figure 7 -.**
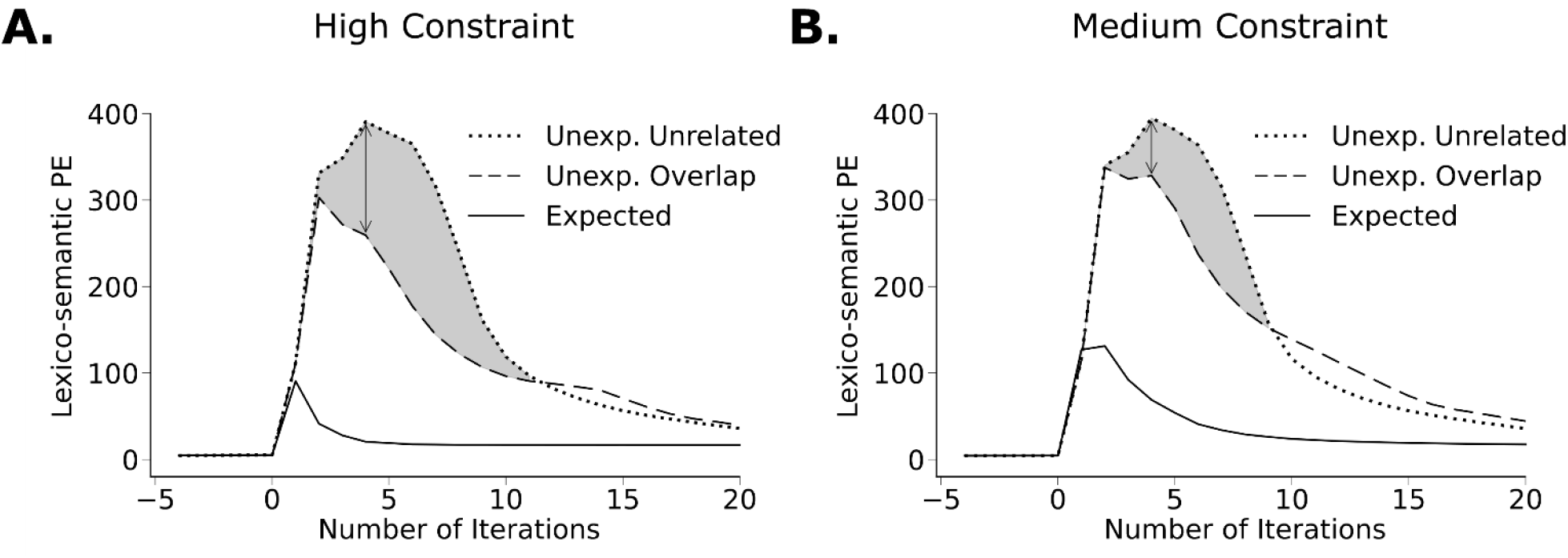
Effects of anticipatory semantic overlap on the time course of lexico-semantic prediction error. A. In the *high constraint* condition (in which the model was pre-activated with 99% probability), lexico-semantic prediction error was largest to the *unexpected unrelated* words (Unexp. Unrelated), smaller to the *unexpected semantically overlapping* words (Unexp. Overlap) and smallest to the *expected* words. **B.** In the *medium constraint* condition (in which the model was pre-activated with 50% probability), lexico-semantic prediction error also decreased across the three conditions. However, as indicated using arrows/shading, in this *medium constraint* condition, the *difference* in prediction error produced by the *unexpected unrelated* and the *unexpected semantically overlapping* words was smaller than this difference in the *high constraint* condition.

The attenuation of lexico-semantic prediction error to the *unexpected semantically overlapping* words arose because the strong top-down prediction of the expected word pre-activated semantic features that were shared with *unexpected semantically overlapping* input, which, in turn, produced reconstructions that suppressed the magnitude of lexico-semantic prediction error produced by these inputs when they were presented.

#### 3.4 Effect of contextual constraint on the anticipatory semantic overlap effect

In addition to showing an effect of anticipatory semantic overlap on the N400 produced by unexpected words in high constraint contexts, Federmeier and Kutas (1999) also showed that this effect interacted with contextual constraint (see also Ito, et al., 2016 for a recent replication using a different set of materials); that is, the degree to which the N400 was reduced in response to the *unexpected semantically overlapping* words (“pines”), relative to the *unexpected unrelated* words (“tulips”), was *greater* in the *high constraint* contexts than in the *medium constraint* contexts. This was taken as evidence that the effect was truly anticipatory: *high constraint* contexts pre-activated the semantic features of the anticipated word to a greater degree than in the *medium constraint* contexts, resulting in greater facilitation on lexically unexpected words that shared these features.

To simulate this interaction between Semantic Overlap and Constraint, we carried out the same simulations as described above, except that instead of assigning the dummy states a probability of 99%, we assigned them a probability of 50%. We then compared the magnitude of prediction error produced by the *unexpected unrelated* and the *unexpected semantically overlapping* inputs across the *high constraint* and *medium constraint* conditions. As shown in Figure 7B, there appeared to be a smaller difference in lexico-semantic prediction error between the *unexpected unrelated* and *unexpected semantically overlapping* words in the *medium constraint* condition (17% reduction) than in the *high constraint* condition (35% reduction). This was confirmed by a linear mixed-effects model that crossed Semantic Overlap (*semantically overlapping*, *unrelated*) with Constraint (*high constraint*, *medium constraint*), which showed that, in addition to a main effect of Semantic Overlap (*b* = -45.45, *t* = -22.83, *p* < .001) and no main effect of Constraint (*p* = .48), there was a significant interaction between Semantic Overlap and Constraint (*b* = -49.82, *t* = -17.69, *p* < .001). This is because the semantic features of the inputs received stronger pre-activation in the *high constraint* (99%) than in the *medium constraint* (50%) condition, leading them to produce more accurate reconstructions that resulted in a greater suppression of the lexico-semantic prediction error to the lexically unexpected input.

#### 3.5 Effect of anticipatory orthographic overlap on words

The N400 is also sensitive to the *orthographic* overlap between a predicted and encountered input (Laszlo & Federmeier, 2009; Ito, Corley, Pickering, Martin & Nieuwland, 2016; DeLong, Chan & Kutas, 2019). For example, Laszlo and Federmeier (2009) presented participants with highly constraining sentence contexts, e.g. “*The genie granted his third and final…*”, followed by critical words that were highly predictable (e.g., *wish*), unexpected (<1% probability) but with several letters in common with the expected word (e.g., *dish*), or unexpected (<1% probability) with no letters in common with the expected word, but matched to the two other conditions on orthographic neighborhood size (e.g. *clam*). The authors observed a graded reduction of the N400 across the three conditions (*wish* < *dish* < *clam*).

To simulate this *anticipatory orthographic overlap* effect, we used 400 words from a set of stimuli that we developed for the purpose of these simulations (see section 3.6 for details). We pre-activated the model with each of these 400 words, assigning the dummy states a probability of 99% and clamping them for 20 iterations. We then presented the model with (a) the same word that was pre-activated (*expected*), (b) a different word that shared three letters (but no semantic features) with the expected word (*unexpected orthographically overlapping*), or (c) a different word that had minimal orthographic overlap (and no semantic overlap) with the expected word (*unexpected unrelated*).

As shown in Figure 8A, consistent with the empirical findings, we saw a graded reduction of lexico-semantic prediction error across the three conditions, with the *unexpected unrelated* words producing a significantly larger prediction error than the *unexpected orthographically overlapping* words (*b =* 134.70; *t* = 115.47, *p* < .001), which, in turn, produced a significantly larger prediction error than the *expected* words (*b =* 119.65; *t* = 97.77, *p* < .001).

**Figure 8 -.**
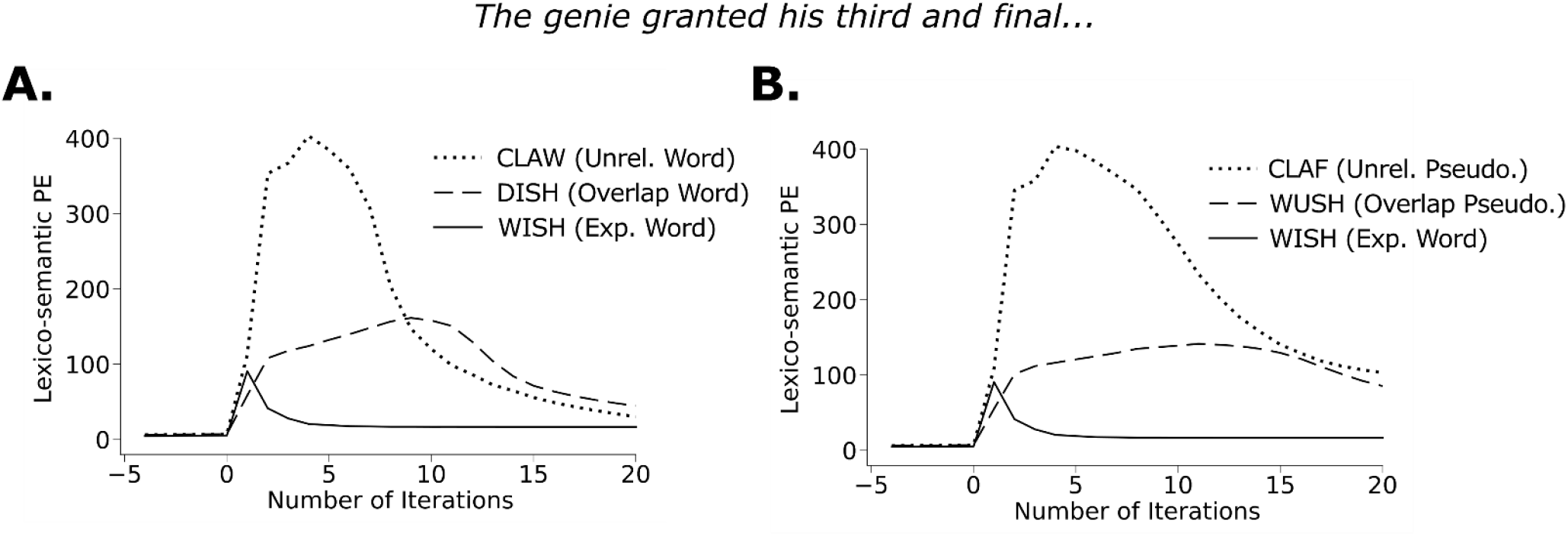
Effect of anticipatory orthographic overlap on the time course of lexico-semantic prediction error. A. Effect of anticipatory orthographic overlap on words. Lexico-semantic prediction error was largest to the *Unexpected unrelated* words (CLAW, Unrel. Word), smaller to the *unexpected orthographically overlapping* words (DISH, Overlap. Word) and smallest to *expected* words (WISH, Exp. Word). **B.** Effect of anticipatory orthographic overlap on pseudowords. Lexico-semantic prediction error was largest to the *Unexpected unrelated* pseudowords (CLAF, Unrel. Pseudo.), smaller to the *unexpected orthographically overlapping* pseudowords (WUSH, Overlap Pseudo.) and smallest to *expected* words (WISH, Exp. Word).

The reason why the *unexpected orthographically overlapping* words (DISH) produced a smaller lexico-semantic prediction error than the *unexpected unrelated* words is because pre-activating the model with the expected word (*wish*) resulted in the generation of orthographic reconstructions (W-**I**-**S**-**H**) that partially suppressed the orthographic prediction error produced by the bottom-up input (D-**I**-**S**-**H**). As a result, less orthographic prediction error flowed up the hierarchy, and so the magnitude of lexico-semantic prediction error was also reduced.

#### 3.6 Effect of anticipatory orthographic overlap on pseudowords

In addition to showing an anticipatory orthographic overlap effect on real words, Laszlo and Federmeier (2009) demonstrated the same effect on pseudowords; that is, the N400 was smaller in response to pseudowords that shared letters with an expected word (e.g. WUSH) than to unrelated pseudowords that were matched on orthographic neighborhood size (e.g. CLAF). To demonstrate this effect, the authors developed a set of stimuli that allowed them to cross Orthographic Overlap (*orthographically overlapping* vs. *unrelated*) and Lexical Status (*word*, *pseudoword*), and match all four conditions on orthographic neighborhood size.

For our simulations, following Laszlo and Federmeier (2009), we developed a set of 400 real word and 400 matched pseudoword stimuli from our 1,579-word lexicon. We began with a set of 400 “base-words”, and, for each of these base words, we constructed a quadruplet of items, thereby setting up a 2 x 2 design that crossed Orthographic Overlap of the critical item with the expected base word (*orthographically overlapping*, *unrelated*) and Lexical Status (*word*, *pseudoword*), while matching items across the four conditions on orthographic neighborhood size. First, for each base-word (WISH), we selected a real word (DISH) and a pseudoword (*WUSH) that overlapped with it in three letter positions, and was matched to it on orthographic neighborhood size (mean ONsize across conditions: 6.71, SD: 3.58). Second, to ensure that each item appeared in both the *orthographically overlapping* and the *unrelated* conditions, we symmetrically paired each base-word (WISH) with another base-word (CLAY) that was of the same orthographic neighborhood size, with the restriction that neither base-word overlapped at any letter position with the other’s word or pseudoword orthographic neighbor (DISH*/*CLAW or *WUSH*/\**CLAF). This yielded 400 items in each of the four conditions: (a) *orthographically overlapping word* (e.g. DISH), (b) *unrelated word* (e.g. CLAW), (c) *orthographically overlapping pseudoword* (e.g. *WUSH), and (d) *unrelated pseudoword* (e.g. *CLAF).

After first pre-activating the model with the base word (WISH, 99% probability) for 20 iterations, we then presented the model with each of the four possible continuations (DISH*, \**WUSH, CLAW*, \**CLAF) in each quadruplet, each for 20 iterations. As shown in Figure 8B, just as for the real word stimuli, *unexpected orthographically overlapping* pseudowords (WUSH) produced a lexico-semantic prediction error that was smaller than that produced by the *unexpected unrelated* pseudowords (*CLAF), but larger than that produced by the *expected* words (WISH). A linear mixed-effects model that crossed Orthographic Overlap (*orthographically overlapping*, *unrelated*) and Lexical Status (*word*, *pseudoword*) confirmed a main effect of Orthographic Overlap (*b* = -87.75, *t* = -119.58, *p* < .001). Indeed, the effect on pseudowords was even larger than that on words (Lexical Status x Orthographic Overlap: *b* = 20.39, *t* = 27.80, *p* < .001; note that Laszlo and Federmeier, 2009 found that the effect of Orthographic Overlap was the same on words and pseudowords). Finally, consistent with the simulations reported in section 1.2, we saw a main effect of Lexical Status (*pseudowords* > *words*, *b =* 13.42, *t =* 18.29, *p* < .001) because the model was unable to settle on a single lexico-semantic state that could explain the bottom-up pseudoword input, resulting in a larger lexico-semantic prediction error overall.

### 4. Interactions between lexical variables and (a) repetition priming and (b) lexical probability

In section 1, we described simulations of the effects of several lexical variables on the N400 produced by words presented in isolation, and in sections 2 and 3, we described simulations of priming and contextual effects, respectively. There is also empirical evidence that the effects of lexical variables on the N400 can be modulated by both repetition priming and lexical probability based on the prior context. For example, in priming paradigms, the effects of frequency and semantic richness/concreteness are reduced on repeated relative to non-repeated targets (Repetition x Frequency: Rugg, 1990; Repetition x Richness/Concreteness: Rabovsky, Sommer, & Abdel Rahman, 2012a; Kounios & Holcomb, 1994). Similarly, during sentence comprehension, the effects of frequency and semantic richness/concreteness are smaller on predictable words than on unpredictable words (Probability x Frequency: Dambacher, Kliegl, Hofmann, & Jacobs, 2006; Probability x Concreteness: Holcomb, Kounios, Anderson, & West, 1999).

Currently, it is less clear how contextual factors influence the magnitude of the orthographic neighborhood effect on the N400. In word lists, Laszlo and Federmeier (2011) described a slightly smaller effect of orthographic neighborhood size on repeated than non-repeated words, although no statistical tests were reported. However, during sentence comprehension, two ERP studies showed that the effect of orthographic neighborhood size was the same on words that were unpredictable because they appeared early in the sentence (producing a larger N400), as on words that were predictable because they appeared later in the sentence (producing a smaller N400; Payne & Federmeier, 2018; Payne, Lee & Federmeier, 2015). This suggests that, in contrast to other lexical variables, the predictability of a word in sentence context may not override the effect of orthographic neighborhood size on the N400. However, no previous study has orthogonally manipulated lexical probability and orthographic neighborhood size in an experimental design.

Here, we simulated interactions between each of our lexical variables — ONsize, Frequency and Richness — with both repetition priming and lexical probability, by re-analyzing data from the simulations described above.

#### 4.1 Interactions between lexical variables and repetition priming

In Figure 9 (top row), we show the effects of ONsize, Frequency and Richness on the magnitude of lexico-semantic prediction error produced by *repeated* versus *non-repeated* target words in our repetition priming simulations. In all three cases, the effects of each of these lexical variables appeared to be smaller on the *repeated* than the *non-repeated* targets. This was confirmed by a linear mixed effects model, which included both main effects (Repetition, ONsize, Frequency and Richness) as well as three interaction terms (Repetition x ONsize, Repetition x Frequency, and Repetition x Richness). This analysis confirmed main effects of Repetition (*b* = -125.52, *t* = -232.81, *p* < .001) and each of the three lexical variables (ONsize: *b* = 15.37, *t* = 28.47, *p* < .001; Frequency: *b* = -1.62, *t* = -3.00, *p* = .003; Richness: *b* = 6.46, *t* = 11.95, *p* < .001). Critically, it also revealed interactions between Repetition and all three lexical variables (Repetition x ONsize: *b* = -15.13, *t* = -28.02, *p* < .001; Repetition x Frequency: *b* = 1.29, *t* = 2.39, *p* = .017; Repetition x Richness: *b* = -3.22, *t* = -5.96, *p* < .001), which, in all cases, were driven by smaller effects on repeated than non-repeated words (ONsize: 99% reduction, Frequency: 89% reduction, Richness: 67% reduction).

**Figure 9 -.**
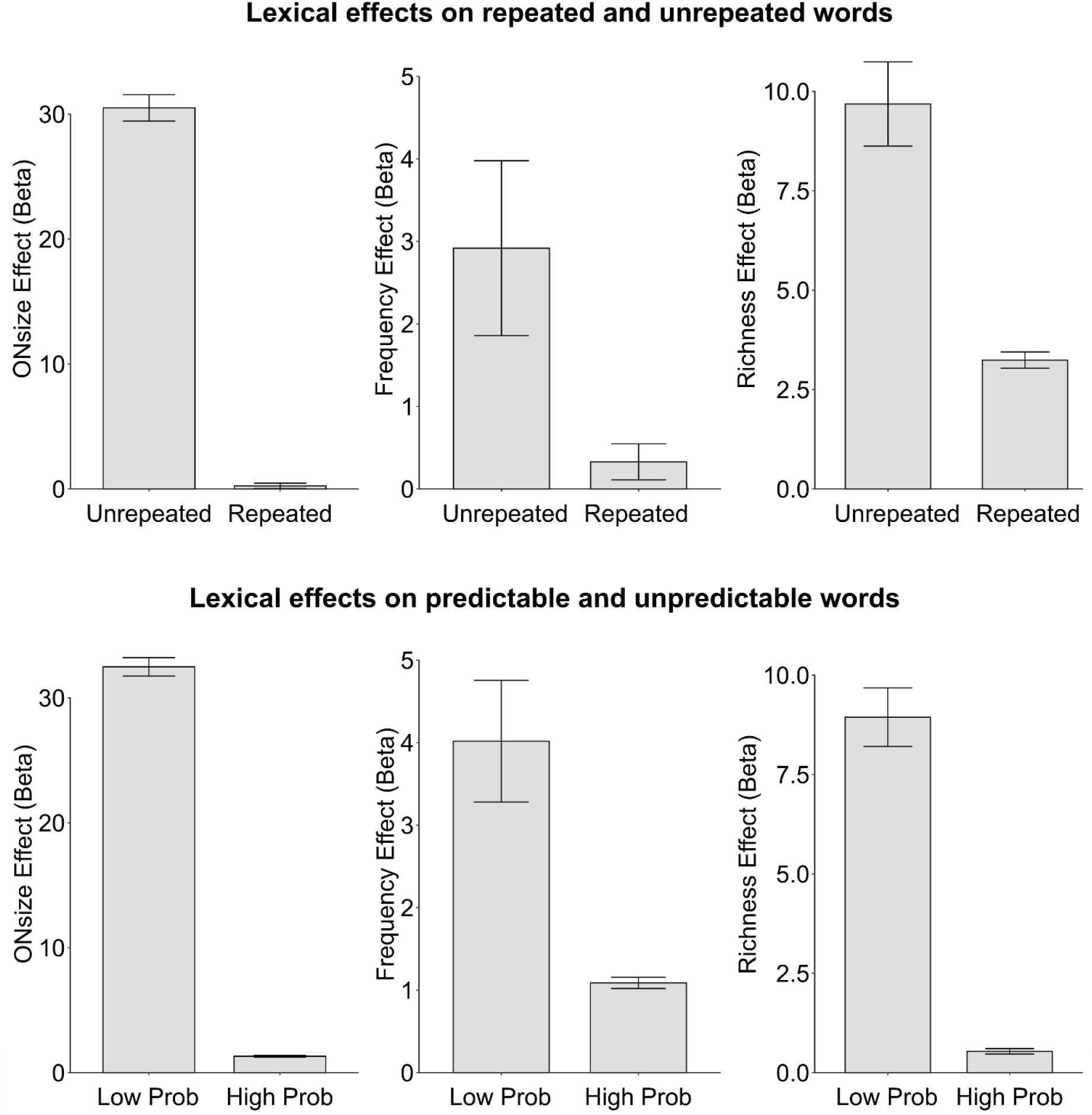
Interaction between lexical variables and repetition priming (top row), and lexical probability (bottom row). In all bar charts, the y-axis shows the average estimate of the slope (i.e., the beta value) obtained by regressing ONsize, Frequency and Richness on the lexico-semantic prediction error. Error bars indicate ±1 standard error of the mean. The effects of all three lexical variables on the magnitude of lexico-semantic prediction error were reduced in the repeated (*vs.* non-repeated) conditions, and in the high (*vs.* low) probability conditions.

#### 4.2 Interactions between lexical variables and lexical probability

In Figure 9 (bottom panel), we show the effects of the same three lexical variables on the magnitude of lexico-semantic prediction error produced by highly expected (99%) and unexpected (0.06%) words. The effects of each of these lexical variables appeared to be smaller on the *expected* than the *unexpected* critical words. Again, this was confirmed by a linear mixed effects model that included all four main effects (Probability, ONsize, Frequency, and Richness) and three interaction terms (Probability x ONsize, Probability x Frequency, and Probability x Richness). In addition to confirming main effects of Probability (*b* = -94.35, *t =* -352.47, *p* < .001) and each of the three lexical variables (ONsize: *b* = 18.79, *t* = 49.58, *p* < .001; Frequency: *b* = -2.73, *t* = -7.19, *p* < .001; Richness: *b* = 5.25, *t* = 13.84, *p* < .001), we found interactions between Probability and each lexical variable (Probability x ONsize: *b* = -11.53, *t =* -43.00, *p* < .001; Probability x Frequency: *b* = 1.08, *t =* 4.03, *p* < .001; Probability x Richness: *b* = -3.11, *t =* -11.58, *p* < .001). In all cases, these interactions arose because the effect of each lexical variable was smaller when the word was predictable than when it was unpredictable (ONsize: 96% reduction, Frequency: 73% reduction, Richness: 94% reduction).

These under-additive interactions receive a straightforward interpretation within our predictive coding framework. Just as for words presented in isolation (see section 1), in the non-repeated and unexpected conditions, higher frequency words produced a smaller lexico-semantic prediction error than lower frequency words because the stronger feedback weights allowed for the generation of more accurate predictions. In the case of repeated or highly predictable targets, however, this default prior was overridden, and so the magnitude of lexico-semantic prediction error was small, regardless of frequency. Similarly, the interactions with Richness arose because the additional unpredicted lexico-semantic information (i.e., lexico-semantic prediction error) carried by words with many (*versus* few) semantic features was reduced when these words were repeated or strongly pre-activated in context. Finally, the reason why the effect of orthographic neighborhood size was reduced in response to repeated/expected target words in our simulations is that accurate reconstructions reduced the magnitude of orthographic prediction error, which, in turn, reduced the magnitude of lexico-semantic prediction error.

## Discussion

We have demonstrated, for the first time, that the basic principles of predictive coding, a general computational theory of brain function, can successfully explain key properties of the N400 response –– a key neural signature of language comprehension. Using a predictive coding model with an architecture and algorithm that were initially developed to simulate a variety of phenomena in vision (Rao & Ballard, 1999; Spratling, 2013; Spratling, 2014), we were able to reproduce a wide range of lexical, priming and contextual effects in the N400 literature. While previous models have successfully simulated some of these effects, the current predictive coding model was able to simulate word-level, priming *and* contextual effects, including several phenomena that have not previously been simulated. It also provides a natural explanation for the temporal dynamics of the N400, and a biologically plausible link to neural activity.

### The predictive coding optimization algorithm explains the rise-and-fall morphology of the N400

Language processing can be understood as probabilistic inference (Chater, Crocker & Pickering, 1998; Narayanan & Jurafsky, 2002; Levy, 2008; Norris, 2006; Kuperberg & Jaeger, 2016). Within this probabilistic generative framework, the brain is assumed to encode an internal generative model that specifies how an underlying higher-level *cause* “generates” observable linguistic data. Inference involves inverting this generative process by using the observed data to estimate the underlying representations (causes) that gave rise to it. Predictive coding is an optimization algorithm that has been proposed to approximate this inferential process in the brain. At each level of the representational hierarchy, “state units” generate top-down predictions (or “reconstructions”) of the hypothetical data at the level below. The difference between the observed data and this prediction is encoded within “error units” as *prediction error*. The predictive coding algorithm functions to minimize this error by using it to modify higher-level states so that they generate more accurate predictions on the next iteration of the algorithm. This process repeats over multiple iterations until the model converges to a set of states whose predictions closely match the observed data, at which point prediction error is minimized.

In our hierarchical predictive coding model, semantic states encode the underlying causes of lexical states, which encode the underlying causes of orthographic inputs. The N400 is operationalized as the total prediction error produced by the model at the lexical and semantic levels as the predictive coding algorithm infers meaning from the orthographic form of incoming words. Thus, prediction error is computed within the model itself, and it plays a crucial role in inference. This distinguishes predictive coding from other computational models that have also operationalized the N400 as “prediction error” (Rabovsky & McRae, 2014; Fitz & Chang, 2019) or as a “change of state” (Rabovsky, Hansen & McClelland, 2018; Brouwer, Crocker, Venhuizen & Hoeks, 2017), but in which these difference values are computed externally by the modeler, play no causal role in comprehension, and remain static over time (see Nour Eddine, Brothers, & Kuperberg, 2022 for discussion).

In predictive coding, the integral role that prediction error plays in inferring higher-level states provides a natural explanation for why the N400 rises and falls over time: the total lexico-semantic prediction error (the N400) produced by the model corresponds to the total neural activity produced by individual lexical and semantic error units, and the rise-and-fall of this error over time emerges from the temporal dynamics of the algorithm (see also Friston, 2005 for discussion in relation to earlier ERP components). Specifically, when new unpredicted orthographic input is presented, lexical and semantic states are updated, resulting in a rise in lexico-semantic prediction error (an increase in the residual difference between the updated states and top-down predictions), explaining the rise in N400 amplitude. Then, as this error is passed up the hierarchy, and the lexical and semantic states converge to stable values, these updated states produce more accurate top-down predictions that suppress the lexico-semantic prediction error, resulting in the fall in N400 amplitude^3^.

### A single measure –– lexico-semantic prediction error –– captures lexical, priming and context <u>effects on the N400</u>

The multiplicity of factors that can affect the amplitude of the N400 has raised questions about whether this ERP component can be understood as a univariate error signal (Federmeier, 2022). Our model tackles this challenge directly. We demonstrate that lexical, priming *and* higher-level contextual effects on the N400, as well as their higher-order interactions, can be captured by a single dependent measure: *lexico-semantic prediction error* — the total activity produced by error units at the lexical and semantic levels as the model processes bottom-up orthographic input.^4^

As noted in the Introduction, previous computational models of the N400 have primarily focused on simulating *either* word-level *or* sentence-level phenomena. Our predictive coding model is unique in that it was able to simulate *both* word-level and higher-level context effects. This is because, similar to classic interactive-activation networks (McClelland & Rumelhart, 1981; McClelland & Elman, 1986), the hierarchical predictive coding architecture allows for a continuous, interactive top-down and bottom-up flow of information across multiple levels of representation (Rao & Ballard, 1999; Mumford, 1992; Friston, 2005) — a crucial feature of online language comprehension (Marslen-Wilson, 1987; Marslen-Wilson, Brown & Tyler, 1988; MacDonald, Pearlmutter & Seidenberg, 1994).

#### Simulations of word-level phenomena, including lexical status

Similar to previous word-level models (Laszlo & Plaut, 2012; Laszlo & Armstrong, 2014; Cheyette & Plaut, 2016; Rabovsky & McRae, 2014), our predictive coding model was able to explain why the amplitude of the N400 is attenuated when words are repeated or semantically primed. It was also able to explain why, when words are presented in isolation, the N400 is sensitive to various lexical characteristics. Notably, this not only included the effects of orthographic neighborhood size on words (B-A-L-L), but also on pseudoword letter strings (*W-E-E-N), which has thus far only been successfully simulated by Laszlo & Plaut (2012). This is because, as originally discussed by Laszlo & Federmeier (2011), the co-activation of orthographic neighbors will result in the activation of their associated semantic features, regardless of whether the eliciting stimulus is a word or a non-word.

In addition, we were also able to simulate for the first time, the larger N400 that is often reported in response to pseudowords, relative to words (cf. Bentin, 1987), even when holding orthographic neighborhood size constant (Holcomb, Grainger & O’Rourke, 2002; Meade, Midgley, Dijkstra & Holcomb, 2018; Braun, Jacobs, Hahne, Ricker, Hofmann & Hutzler, 2006; but see Laszlo & Federmeier, 2011). The reason for this is that a pseudoword string input (e.g., *W-E-E-N) activated multiple lexical units in parallel (e.g. *weed*, *teen*, *wren* etc.). However, in contrast with real word inputs, the model failed to converge on a single representation, resulting in a failure to fully suppress lexico-semantic prediction error.

#### Simulations of broader contextual effects through top-down pre-activation

Similar to previous sentence-level models, our predictive coding model was able to simulate several effects of broader context on the N400. These included the graded effects of lexical predictability (Kutas & Hillyard, 1984; DeLong et al., 2005; simulated by Rabovsky, Hansen & McClelland, 2018; Fitz & Chang, 2019; see also Brouwer et al., 2017), the null effect of contextual constraint over and above lexical predictability (Kutas & Hillyard, 1984; Federmeier et al., 2007; simulated by Rabovsky, Hansen & McClelland, 2018; Fitz & Chang, 2019), and the attenuation of the N400 to unexpected words that semantically overlap with expected words (Federmeier & Kutas, 1999, simulated by Rabovsky, Hansen & McClelland, 2018).

Notably, however, we took a different approach to modeling these higher-level context effects from that taken by these previous models. In many psycholinguistic theories of predictive processing, it is assumed that a higher-level interpretation of the prior context is used to *pre-activate* lower-level lexico-semantic states *prior* to new bottom-up input becoming available (Federmeier, 2007; DeLong, Urbach & Kutas, 2005; Kuperberg & Jaeger, 2016, Section 3, pp. 39-45 for a detailed discussion; for direct neural evidence of lower-level pre-activation, see Wang, Hagoort, & Jensen, 2018; Hubbard & Federmeier, 2021). However, in previous models of the N400, higher-level representations did not actually pre-activate expected information at a lower lexico-semantic level *before* the onset of the bottom-up input. For example, in the model by Brouwer et al. (2017), context only exerted an influence on an intermediate semantic layer *after* the expected input appeared, and in the model by Rabovsky, Hansen & McClelland (2018), the semantics of predictable future inputs and the prior context were encoded within the same layer that was used to simulate the N400.

In our simulations, we directly implemented *top-down* pre-activation by providing inputs to the highest layer of the model’s hierarchy (although, as noted below under Future Directions, this “dummy layer” did not directly encode higher-level representations, but rather a probability-like distribution over words in the model’s lexicon). This information was then allowed to filter down the hierarchy and pre-activate state units at the lower semantic and lexical layers, prior to the presentation of new bottom-up orthographic input. This type of top-down pre-activation fulfills two important computational functions during natural language processing: First, it offers a *head-start* in processing information at lower levels, increasing the speed and efficiency of accessing the meaning of incoming words as they unfold rapidly in real time. Second, information from higher levels provides a top-down *bias* that “fills in” lower-level state representations, making the system robust to perceptual noise or ambiguous inputs (see Mumford, 1992 for discussion) — a form of “pattern completion” in the temporal domain (cf. Barlow, 1998; Falandays, Nguyen & Spivey, 2021; Huettig, Audring & Jackendoff, 2022), but at multiple levels of the representational hierarchy.

This top-down pre-activation mechanism allowed us to carry out simulations of certain contextual effects that previous models had not simulated. First, we were able to simulate the effect of contextual constraint on the magnitude of the *anticipatory semantic overlap effect* (Federmeier & Kutas, 1999). The semantic overlap effect refers to the reduction of the N400 to incoming words that, although lexically unexpected, are nonetheless semantically related to the expected continuation. In an important study, Federmeier & Kutas (1999) found that this effect was modulated by contextual constraint, even when the local semantic relationships between the incoming word and the immediately preceding words were held constant, and even though the input was implausible. This led the authors to propose that the semantic overlap effect was driven by the *top-down pre-activation* of semantic features of the expected upcoming word by the broader context. On this account, the more strongly the broader context constrains for a specific upcoming word, the more strongly its semantic features are pre-activated, and so the more facilitation an incoming word that shares these semantic features will receive. The present simulations directly support this interpretation: the more top-down pre-activation the model was given for a specific upcoming word, the smaller the lexico-semantic prediction error produced by an incoming word that was unexpected but semantically related to the expected continuation.

Second, our top-down pre-activation mechanism allowed us to simulate, for the first time, the anticipatory *orthographic* overlap effect — the smaller N400 elicited by unexpected inputs that overlap orthographically with an expected continuation (e.g., to the word, “dish”, following the context, “The genie granted his third and final…”; Laszlo and Federmeier, 2009; Ito, Corley, Pickering, Martin & Nieuwland, 2016; DeLong, Chan & Kutas, 2019). Moreover, mirroring Laszlo and Federmeier’s (2009) empirical findings, we were also able to simulate the smaller N400 produced by *pseudoword* letter strings that overlapped with expected continuations (e.g., W-U-S-H). Predictive coding offers a natural explanation for these anticipatory orthographic overlap effects. By providing the model with top-down pre-activation, the lexical state unit that corresponded to the expected word (e.g., *wish*) generated orthographic reconstructions of the letters, W-I-S-H. When the unexpected orthographic inputs were presented (D-I-S-H or W-U-S-H) the reconstruction of letters that overlapped with these inputs (_-I-S-H or W-_-S-H) suppressed the generation of orthographic prediction error. This small orthographic prediction error, in turn, drove only small changes in state at the lexical and semantic levels, resulting in minimal prediction error at the lexical and semantic levels.

Finally, the interactive nature of the predictive coding architecture, with its free flow of top-down and bottom-up information across hierarchical levels, allowed us, for the first time, to model the *interactions* between lexical probability and lexical variables on the N400. Mirroring the empirical findings, we showed that the effects of frequency and semantic richness on the magnitude of lexico-semantic prediction error were attenuated, not only to repeated targets in priming paradigms (an interaction that was successfully simulated by previous models, see Rabovsky & McRae, 2014; Cheyette & Plaut, 2017), but also when they were contextually predictable (empirical findings for Predictability x Frequency, see Dambacher, Kliegl, Hofmann, & Jacobs, 2006, and for Predictability x Richness, see Holcomb, Kounios, Anderson, & West, 1999). In our simulations, this is because the pre-activation of the target state units generated reconstructions that rapidly suppressed the production of lexico-semantic prediction error in response to expected inputs, regardless of their lexical characteristics.

Of note, our simulations showed that repetition priming and top-down lexical probability interacted not only with frequency and semantic richness, but also with orthographic neighborhood size; that is, the effect of orthographic neighborhood size on the magnitude of lexico-semantic prediction error was smaller when the input was repeated or highly predictable. In contrast to the effects of frequency and semantic richness, previous experimental studies have not directly compared the effects of orthographic neighborhood size on the same items in highly predictable *versus* non-predictable contexts. However, there is some evidence that this effect may be more impervious to the effects of context than the effects of other lexical factors. For example, Payne et al. (2015, 2018) found that, during sentence comprehension, the effect of orthographic neighborhood size on the N400 was just as large to more predictable words that appeared later in sentences than to less predictable words that appeared earlier in sentences. It will therefore be important for future empirical studies and simulations to further explore this interaction (see Future Directions).

### Implications

By showing that the N400 evoked response can be understood as the production of lexico-semantic prediction error in a predictive coding framework, this work helps explain and brings together multiple strands of the prior literature.

First, our finding that the effects of *both* bottom-up priming and top-down lexical predictability on the N400 can be simulated as prediction error generated at the *lexico-semantic* level of representation explains why both these effects localize to regions of the left temporal cortex that are known to support lexico-semantic processing (effects of semantic priming: Nobre & McCarthy, 1995; Lau, Weber, Gramfort, Hämäläinen & Kuperberg; Lau, Gramfort, Hämäläinen & Kuperberg, 2013; effects of lexical probability in plausible sentences: Wang et al., 2022). In both cases, this is because state units are activated prior to the onset of the target – either due to lingering bottom-up activation in the case of priming, or due to top-down pre-activation when a word is predictable in context – and so the magnitude of lexico-semantic prediction will be suppressed when the bottom-up input appears.

Second, our findings directly bridge the large existing N400 literature to previous MEG and fMRI work in speech perception (Blank & Davis, 2016; Sohoglu & Davis, 2020), and visual word recognition (Price & Devlin, 2011), which have interpreted the larger neural response to unexpected relative to expected inputs to prediction error that is generated at lower levels of language hierarchy.

Third, and most generally, our findings link the N400 to the large body of research in predictive coding across non-linguistic perceptual and cognitive domains (Clark, 2013; Spratling, 2016b). Notably, the basic structure and connections of our predictive coding architecture, as well as the steps of the predictive coding algorithm, were based directly on models that were originally developed to explain low-level visual phenomena (Rao & Ballard, 1999; Spratling, 2012, 2013, 2014). The fact that our model was able to simulate such a wide range of effects on the N400 is therefore consistent with the idea that a common canonical circuit motif (cf. Douglas, Martin & Whitteridge, 1989) may implement predictive coding in the brain across multiple domains of perception and cognition (Bastos et al., 2012; Aitchison & Lengyel, 2017), including language comprehension.

### Limitations and future directions

Our model has several limitations, opening up many potential avenues for future research. One important goal will be to expand the representations at the highest level of the hierarchy. In the current version of the model, state units within the highest dummy layer directly encoded the probabilities of upcoming words. Our contextual simulations involved providing top-down inputs directly to this layer and allowing information to propagate down the hierarchy before the bottom-up input was presented. In reality, however, these probabilities would be based on higher-level event-level representations that stretch over a longer time scale and that are inferred incrementally from the full sequence of linguistic inputs (Hare, Jones, Thomson, Kelly, & McRae, 2009; McRae & Matsuki, 2009). Incorporating a high-level *event layer* would allow us to model certain event-level phenomena that previous models were able to simulate, such as the effects of semantic attraction on unexpected words (Kuperberg, Sitnikova, Caplan & Holcomb, 2003; simulated by Brouwer et al., 2017, and by Rabovsky, Hansen & McClelland, 2018), and the effects of word position on the N400 (Van Petten & Kutas, 1990, 1992; simulated by Rabovsky, Hansen & McClelland 2018).

Ultimately, for any model to explain how the brain is able to *deeply* comprehend language, the event layer would also need to receive inputs from real-world knowledge, stored within long-term memory (see Franklin, Norman, Ranganath, Zacks, & Gershman, 2020 for one modeling approach). This would allow future models to simulate the effects of real-world plausibility on the N400 (cf. Hagoort, Hald, Bastiaansen & Petersson, 2004). Based on recent MEG findings, we have suggested that the larger N400 evoked by unexpected implausible (versus plausible) inputs (Nieuwland et al., 2020; Kuperberg, Brothers, & Wlotko, 2020) may reflect prediction error that is generated not only at the lexico-semantic level, but also at the higher event-level when a newly-inferred event cannot be explained by still higher-level predictions based on real-world knowledge (Wang et al. 2022; and see Rao & Ballard, 1999 for discussion in the visual system).

In addition, allowing the event layer to receive inputs from longer-term memory would allow any event-level representations inferred from the linguistic input to be further informed by relevant event “schema” (cf. Rumelhart & Ortony, 1977; Schank & Abelson, 1977; Minsky, 1975), which is necessary to build a “situation model” that links events along causal, temporal and spatial dimensions (Van Dijk & Kintsch, 1983; Zwaan & Radvansky, 1998). This would, in turn, allow future models to simulate two distinct later positive-going components that extend beyond the N400 time-window (between 600-1000ms), and that are only produced when comprehenders are engaged in building a situation model (Brothers, Wlotko, Warnke & Kuperberg, 2020). Two previous computational models have attempted to simulate late positivities (Brouwer et al., 2017; Fitz & Chang, 2019), but they each have several limitations (for discussion, see Nour Eddine, Brothers, & Kuperberg, 2022). For example, neither model distinguishes between two distinct types of late positivities (Van Petten & Luka, 2012; Kuperberg, Brothers & Wlotko, 2020) –– a *late frontal positivity* (Federmeier et al., 2007) that is produced when an unexpected inputs triggers a *successful* update of the situation model, resulting in a plausible interpretation (Kuperberg, Brothers & Wlotko, 2020; Brothers, Wlotko, Warnke & Kuperberg, 2020), and a *late posterior positivity/P600* (Kuperberg, 2007; van de Meerendonk, Kolk, Chwilla & Vissers, 2009) that is typically produced when a linguistic error (e.g. a semantic anomaly) *cannot* immediately be incorporated into the situation model, triggering reprocessing of the bottom-up input (Kuperberg, Brothers & Wlotko, 2020; Brothers, Wlotko, Warnke & Kuperberg, 2020; Wang et al., 2022; Kuperberg, Alexander, & Brothers, under review).

In addition to expanding the model’s hierarchy upwards, it may also be possible to expand the lower levels of the hierarchy so as to include intermediate sub-lexical layers that encode bigrams and trigrams (see Grainger, 2008; Dehaene, Cohen, Sigman, & Vinckier, 2005) –– an orthographic coding scheme that is more realistic than the slot-based approach used in the present study. This would still allow us to simulate the effects of orthographic neighborhood size on the N400. However, this simulated effect may be more resistant to the effects of lexical predictability, in keeping with the empirical data (Payne & Federmeier, 2018; Payne, Lee & Federmeier, 2015).^5^ Moreover, adding these intermediate sub-lexical layers would also allow us to simulate earlier language-related ERP components observed between 150-300ms (see Grainger & Holcomb, 2009), which have been linked to the generation of prediction error at lower levels of the language hierarchy (Price & Devlin, 2011).

Another exciting direction for future work will be to determine whether predictive coding is able to simulate the effects of lexical, priming and contextual factors on *behavior*, i.e., the speed and accuracy of word identification (see Rayner, 1998; Rastle, 2016 for reviews). In the present study, we focused on simulating the N400 ERP response, which we operationalized as the total activity produced at any given time by error units at the lexical and semantic levels (i.e., lexico-semantic prediction error). As we have discussed, in predictive coding, prediction error produced at each level of the linguistic hierarchy plays a critical role in updating state units at the level above. Therefore, at the same time as error units are producing transient prediction error (and an evoked response) in response to unpredicted information, state units are converging on a more accurate estimate of the underlying cause of the input, enabling more accurate top-down predictions (see Wang, Nour Eddine, Brothers, Jensen & Kuperberg, under review, for recent MEG evidence that functionally distinct state and error units are differentially activated by expected and unexpected inputs within the N400 time window). The improved estimate encoded within the state units may play an important role in guiding decision-making during behavioral tasks. In future work, we hope to exploit this aspect of the predictive coding architecture to simulate a wide range of findings in the behavioral literature, and better understand their relationship with the N400.

Finally, a key focus for future research will be to examine the relationship between prediction error, comprehension and learning within the predictive coding framework. The close link between prediction error and learning is well documented in both non-linguistic (Rescorla & Wagner, 1972; Rumelhart, Hinton & Williams, 1986) and linguistic domains (e.g., Elman, 1990; Chang, Dell & Bock, 2006). This link has also been emphasized in previous models of the N400 in which prediction error (Fitz & Chang, 2019; Rabovsky & McRae, 2014) or changes-in-state (Rabovsky, Hansen & McClelland 2018) were computed for the purpose of downstream learning through backpropagation.

As noted earlier, in these previous models, prediction error was computed outside the model’s architecture and played no functional role in comprehension. This contrasts with predictive coding where error is computed locally within each layer for the purpose of inference. This, however, doesn’t imply that the prediction error computed during predictive coding isn’t *also* used for downstream learning. In the current implementation of our model, we made the simplifying assumption that the connection weights between the layers were fixed, and so prediction error played no role in learning. However, this is not an inherent limitation of the predictive coding framework. In fact, predictive coding offers a biologically plausible algorithm for instantiating longer-term learning without introducing any new elements into the architecture (Rao and Ballard, 1999; Whittington & Bogacz, 2019; see Nour Eddine, Brothers, & Kuperberg, 2022 for discussion). Indeed, it has been shown that under certain theoretical assumptions, the prediction error computed during predictive coding converges to the learning signal used for backpropagation (Whittington & Bogacz, 2017; Millidge, Tschantz & Buckley, 2020; Song, Lukasiewicz, Xu, & Bogacz, 2020). Under this scenario, the model would alternate between short-term inference (fixing the connection weights and updating the state units) and longer-term learning (fixing the state units and updating the weights). This would capture the idea that language processing and language learning/adaptation are closely intertwined throughout the lifespan (see Elman, 1990; Kleinschmidt & Jaeger, 2015; Dell & Chang, 2014).

## Conclusion

Predictive coding offers a simple, interpretable and biologically plausible framework to make sense of prediction in language comprehension. Our simulations show that lexico-semantic prediction error within this framework shares a remarkable range of features with the N400 event-related component, from its time course, to its sensitivity to top-down and bottom-up variables, and even interactions between these variables. By mapping the N400 on to a distinct element within a hierarchical generative modeling framework (lexico-semantic prediction error), we situate this key neural component within the broader context of predictive coding research. This paves the way towards understanding how specific disruptions of predictive coding might give rise to the pathological neural responses observed during language processing in neurodevelopmental disorders such as schizophrenia (Brown & Kuperberg, 2015; Fletcher & Frith, 2009).

## Acknowledgments

This work was funded by the National Institute of Child Health and Human Development (R01MHD082527) to G.R.K. We thank Arim Choi Perrachione for her help with the figures. We also thank Jeff Stibel for his support of Drs. Kuperberg and Wang.

## Appendix A. Supplementary data

The supplementary materials include a more technical description of the predictive coding algorithm, weight matrices, hyperparameters, frequency implementation and a derivation of the state update step of the algorithm; they also include a figure showing the localization of the semantic richness effect to the semantic level. Supplementary materials can be accessed at the following link: https://projects.iq.harvard.edu/sites/projects.iq.harvard.edu/files/kuperberglab/files/noureddine_predictivecoding_suppl_materials.zip

## Supplementary Materials: A Predictive Coding Model of the N400

### Samer Nour Eddine, Trevor Brothers, Lin Wang, Michael Spratling, Gina Kuperberg

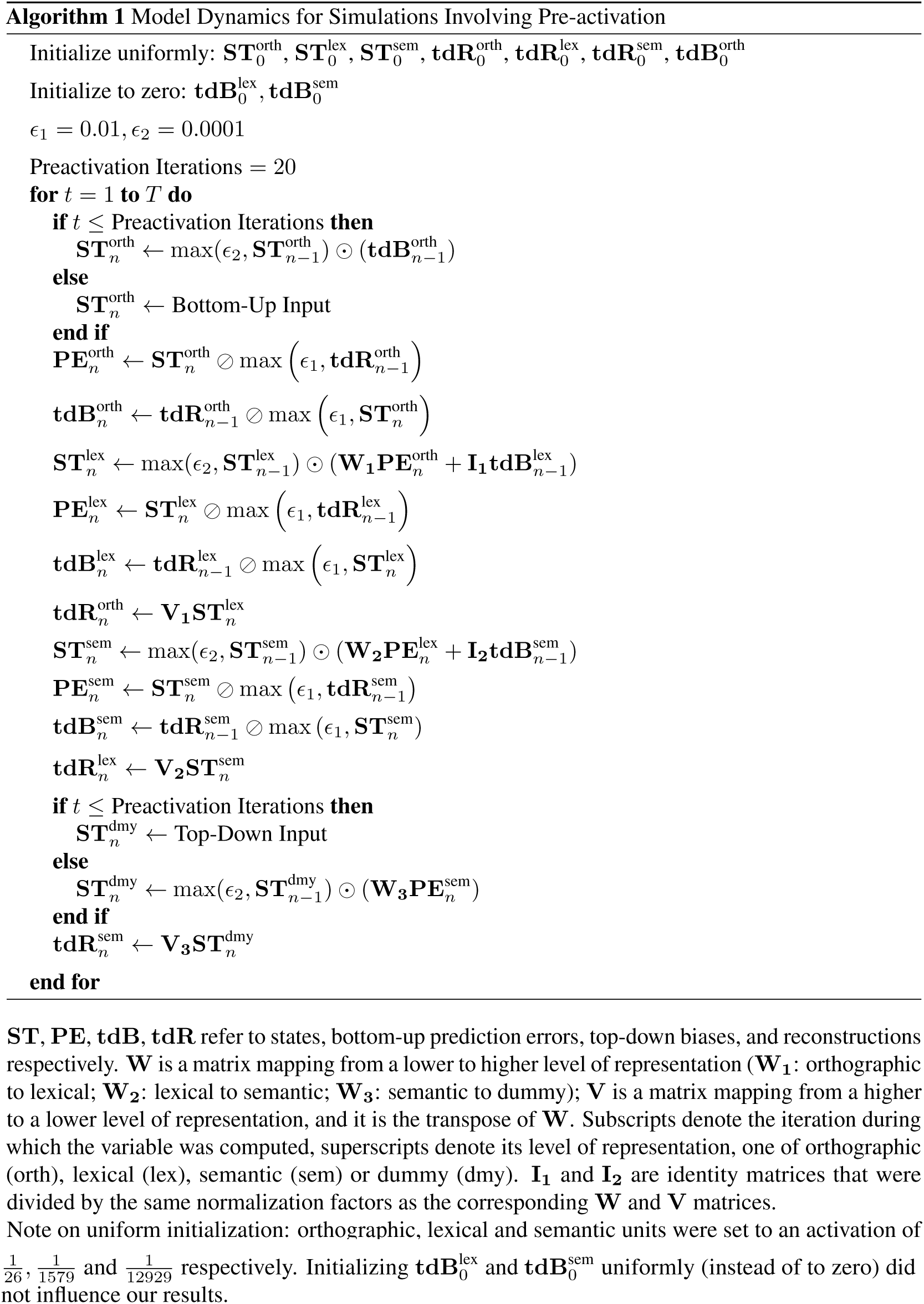

## 1 Weight Matrices

For the orthographic-lexical mappings, we define a pair of matrices V_1_ and W_1_ that are tranposes of one another. Each column of V_1_ encodes the spelling of a particular word. For example, one column encodes the orthographic sequence L-I-O-N, another column encodes the sequence F-R-O-G, and so on for all the orthographic sequences that the model has a lexical representation for. We defined these weight matrix based on known spellings of 1579, four-letter words. For example, consider the column that corresponds to the word *corn*. The first 26 entries in the column correspond to the alphabetical order of the 26 letters of the English alphabet, and can indicate that “C” occupies the first position in *corn* by assigning the 3rd entry the value of 1 and the other 25 entries a value of 0. In the same way, the second 26 entries indicate that “O” occupies the second position, and so on.

For the lexical-semantic mappings, we define a pair of matrices V_2_ and W_2_. Every column in this matrix represents the set of words that have the corresponding semantic feature. For example, one column might track the (animal) semantic feature, and can indicate that *lion* and *frog* have this feature by assigning the column entries corresponding to them a value of 1, and that *lime* and *corn* do not by assigning them a value of 0.

Finally, the semantic-dummy mappings V_3_ and W_3_ were rescaled transposes of W_2_ and V_2_, respectively, and carried out approximately the same mapping role (e.g., similar to W_2_, which projected error from lexical to semantic space, V_3_ projects a *reconstruction* from dummy (which has the same dimensionality as lexical space) to semantic space).

Note that in our implementation, we scaled all V (W) matrices columnwise (rowwise) by a factor that was dictated by the optimization objective (see *Optimization Algorithm* below for details.).

## 2 Hyperparameters

### 2.1 Epsilons

In order to prevent multiplication by zero (which prevents the state from getting updated) and division by zero, we defined two small positive scalars, *E*_1_ and *E*_2_, to be the minimum values the model would divide by or multiply with respectively. These were set to values that allowed the model to converge to a stable state (that approximately minimized a predetermined cost function) as smoothly as possible. We found that no choice of epsilon changed the qualitative patterns that we report in our results, although some choices of epsilon (e.g., *E*_1_ = 0.1) eliminated the cloze probability effect, by capping the prediction error and bias terms at relatively low values and essentially interfering with our manipulation.

### 2.2 Number of Pre-activation Iterations

We found that clamping a top-down input for a reasonable number of iterations - anywhere between 5 and 20 - led to the same results qualitatively. We chose the number 20 in order to match the number of iterations in the post-stimulus window.

## 3 Implementation of Frequency

We start with frequency-naive feedback matrices V_1_, V_2_, V_3_ that map from a higher to a lower level of representation. Frequency information was encoded in the feedback matrices by adding a positive frequency score to each non-zero entry in the frequency-naive feedback matrix. Prior to adding the frequency score, the non-zero entries had the following ranges and means:

Matrix: mean (minimum - maximum)

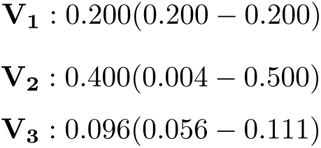

The frequency score was determined by scaling and shifting the log frequency of the word (obtained from Brysbaert & New, 2009) so that it falls into the range (0.0, 0.1):

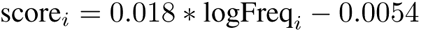

Frequency scores had a mean of 0.041, and ranged from 0.0 to 0.1. Note that each word was associated with (i) a particular frequency score and (ii) a set of non-zero entries in each V matrix. Thus we added each word’s frequency score to all and only the non-zero entries that corresponded to it.

After adding the frequency score, the non-zero V entries had the following ranges and means:

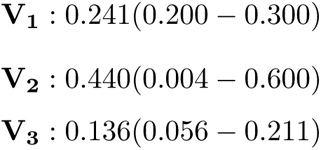

## 4 Optimization Algorithm

Here we derive the update rule for the lexical state units, but an analogous derivation can be applied in the general case. The optimization goal is to find the value for the lexical state units that would minimize two divergence measures. We are interested in finding 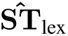 below:

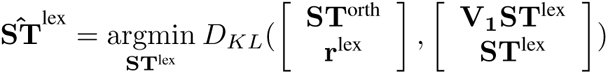

Where 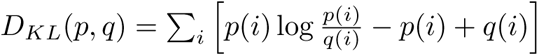, the generalized KL divergence. To simplify, we can rewrite the above as follows:

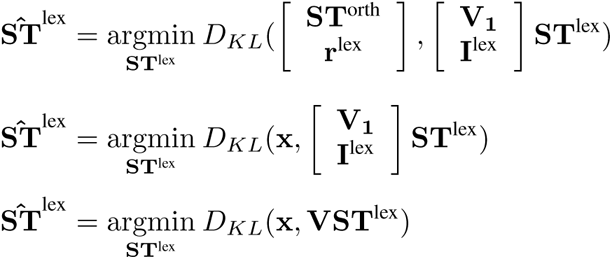

Where I^lex^ is the identity matrix (dimensionality 1579 × 1579); V is the vertical concatenation of the matrix V_1_ (dimensionality orth × lex) and I^lex^; and x is the vertical concatenation of ST^orth^ and r^lex^.

We chose to achieve the optimization goal with gradient descent. The gradient of *D_KL_* with respect to ST^lex^ is the following^1^:

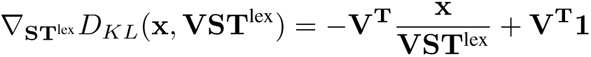

Where V^T^ is the transpose of V; and 1 is a column vector consisting exclusively of ones. By carefully choosing the learning rate (i.e., we put the negative terms of the gradient in the numerator of the update factor, and the positive terms in the denominator of the update factor), the additive update rule of gradient descent simplifies into the following multiplicative rule:

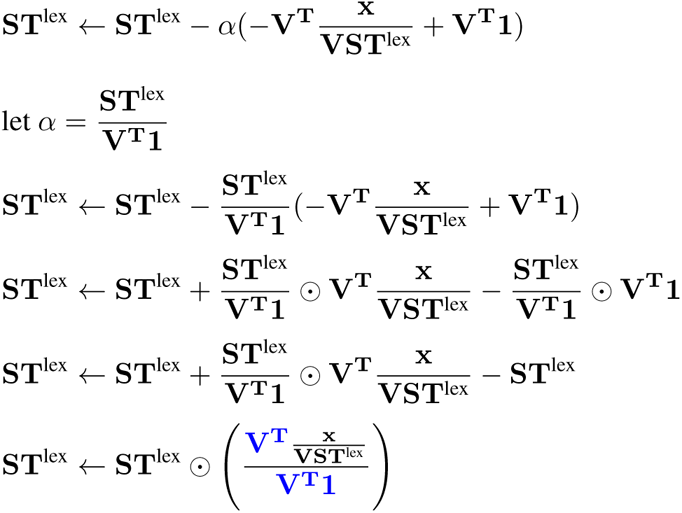

Together, the terms in blue represent a “normalized” matrix **V^T^** where each element in a given row is divided by the sum of the terms in that row. Because each row is scaled separately, we can put all of these scaling factors into a vector *λ*.

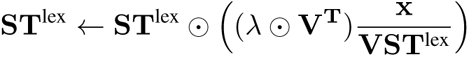

To make the concatenation of 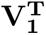 and the identity matrix explicit, we rewrite 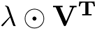 as follows:

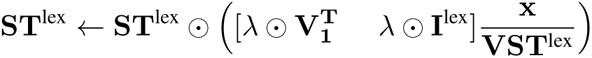

Similarly, we rewrite the 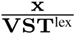 term to make the concatenations explicit:

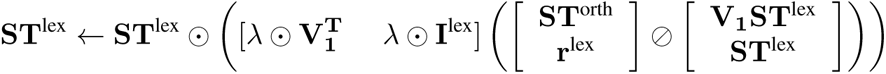

Note that 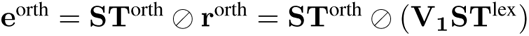, and that 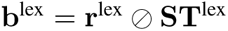. Therefore, we can rewrite the above as follows:

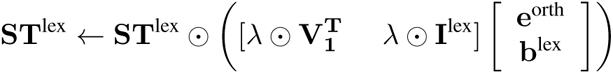

Let 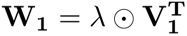, and let 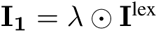. Now we have:

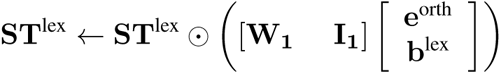

Which simplifies to:

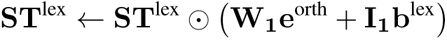

The rest of the update rules can be derived in an analogous manner.

## 5 Richness effect localizes to the Semantic level

**Supplementary Figure 1:**
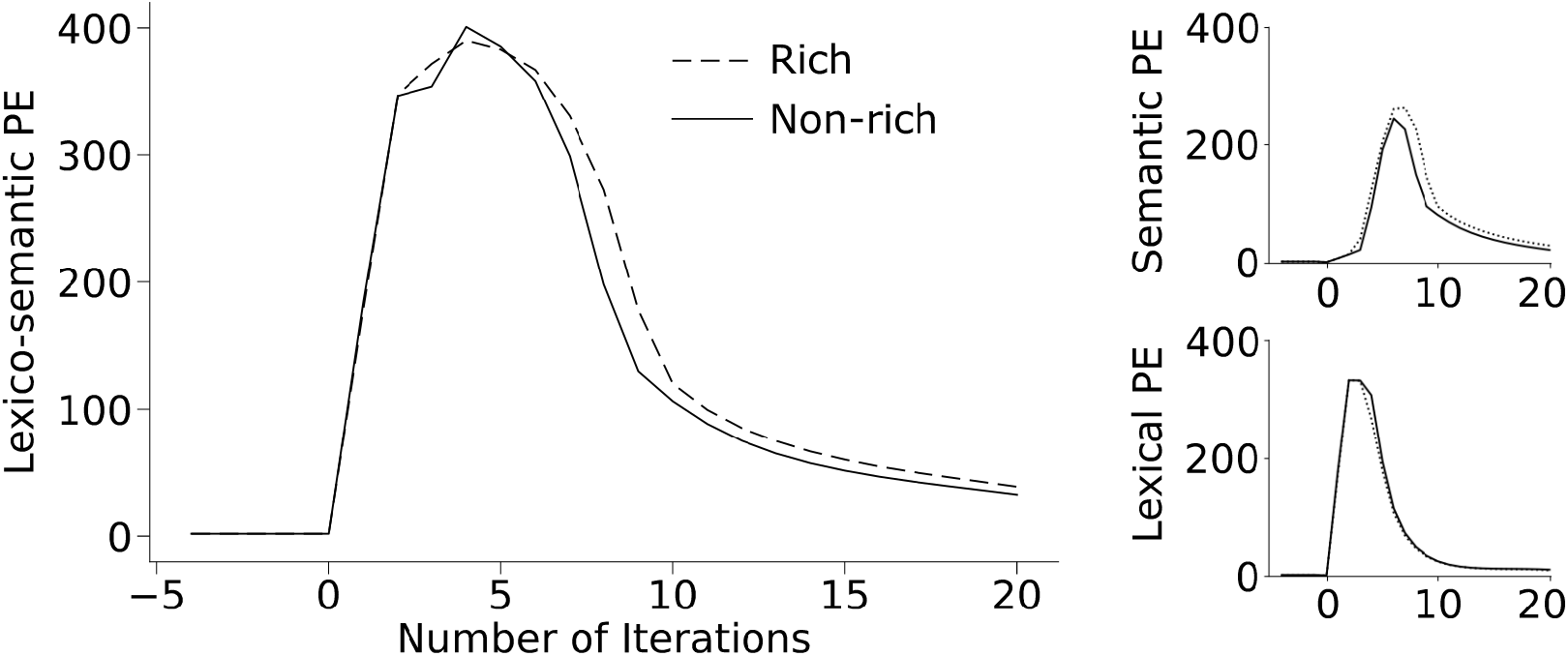
The Richness effect localizes to the Semantic level. Rich items elicit a significantly larger lexico-semantic prediction error than non-rich items (left panel). However, separating the total lexico-semantic prediction error into its lexical and semantic components reveals that the richness effect occurs exclusively at the semantic level (top-right), not at the lexical level (bottom-right).

## 6 References

Brysbaert, M., New, B. (2009). Moving beyond Kučera and Francis: A critical evaluation of current word frequency norms and the introduction of a new and improved word frequency measure for American English. Behavior research methods, 41(4), 977-990.

## Footnotes

1 As we elaborate further in the Discussion section, in predictive coding, prediction error can also, in principle, be used for downstream learning (Whittington & Bogacz, 2017; Millidge, Tschantz & Buckley, 2020; Song, Lukasiewicz, Xu, & Bogacz, 2020).

2 The scalp distribution of the semantic richness/concreteness effect on the N400 is more frontal than the classic centroparietal N400 effect. It has been hypothesized that this is because the effect originates primarily at the level of semantic features, rather at the lexical level that links these features to linguistic form (e.g. Kounios & Holcomb, 1994). Consistent with this account, an exploratory analysis indeed suggested that the effect of richness stemmed primarily from prediction error produced at the semantic rather than the lexical level (see Supplementary Materials).

3 This account of the N400’s time course is distinct from the one discussed in relation to the Semantic Activation model — the only other model that simulated the morphology of the N400 (Laszlo & Plaut, 2012; Laszlo & Armstrong, 2014; Cheyette & Plaut, 2017; see Nour Eddine, Brothers, & Kuperberg, 2022 for discussion).

4 The additional production of prediction error at a higher event-level may contribute to additional effects of *plausibility* on the N400 (see Wang et al., 2022 and Future Directions for discussion).

5 Using bigrams and trigrams enables a more efficient orthographic coding scheme, allowing the model to represent words that have smaller orthographic neighborhood sizes (e.g., *kiwi*) with fewer units. For example, fewer bigrams are required to uniquely identify the lexical item *kiwi* than the item *core* which has a larger orthographic neighborhood size. Therefore, under this scheme, reconstructing the word-form of KIWI correctly would require fewer units compared to reconstructing the word-form of CORE. Because the resulting orthographic neighborhood size effect would be driven by a difference in word-form *encoding*, it should be relatively invariant to top-down factors like lexical predictability.

1 Derived here: https://www.jjburred.com/research/pdf/jjburrednmfupdates.pdf

